# ProtAnno, an Automated Cell Type Annotation Tool for Single Cell Proteomics Data that integrates information from Multiple Reference Sources

**DOI:** 10.1101/2021.09.13.460162

**Authors:** Wenxuan Deng, Biqing Zhu, Seyoung Park, Tomokazu S. Sumida, Avraham Unterman, David Hafler, Charles S. Dela Cruz, Naftali Kaminski, Carrie L. Lucas, Hongyu Zhao

## Abstract

Compared with sequencing-based global genomic profiling, cytometry labels targeted surface markers on millions of cells in parallel either by conjugated rare earth metal particles or Unique Molecular Identifier (UMI) barcodes. Correct annotation of these cells to specific cell types is a key step in the analysis of these data. However, there is no computational tool that automatically annotates single cell proteomics data for cell type inference. In this manuscript, we propose an automated single cell **prot**eomics data **anno**tation approach called **ProtAnno** to facilitate cell type assignments without laborious manual gating. ProtAnno is designed to incorporate information from annotated single cell RNA-seq (scRNA-seq), CITE-seq, and prior data knowledge (which can be imprecise) on biomarkers for different cell types. We have performed extensive simulations to demonstrate the accuracy and robustness of ProtAnno. For several single cell proteomics datasets that have been manually labeled, ProtAnno was able to correctly label most single cells. In summary, ProtAnno offers an accurate and robust tool to automate cell type annotations for large single cell proteomics datasets, and the analysis of such annotated cell types can offer valuable biological insights.

## Introduction

Recent years have seen the developments of many single cell platforms(Eberwine et al. 2014) that have enabled researchers to collect high throughput -omics profiles at the individual cell level, including genomics(Dong et al. 2017; Nagano et al. 2013), transcriptomics(Hwang, Lee, and Bang 2018), proteomics(Labib and Kelley 2020), and epigenomics(Fang et al. 2021). These data can reveal biological heterogeneity across different biological conditions. They offer a direct approach to studying cell type compositions and functional cell states. The most well-developed platforms are scRNA-seq for transcriptomics and flow cytometry (CyTOF)(Bandura et al. 2009; Spitzer and Nolan 2016; Ornatsky et al. 2010) for proteomics. Notably, the rise of scRNA-seq has generated rich data and resources on single cell transcriptomics(Regev et al. 2017; Lindeboom, Regev, and Teichmann 2021).

In addition to collecting single cell data for one specific-oimcs data type, it is possible to collect multi-omics data simultaneously at the single cell level, e.g., CITE-seq(Stoeckius et al. 2017) that measures mRNA and antibody counts simultaneously by UMI barcode. These data can characterize the cellular relationship between transcript and cell surface marker abundance. Methods have been developed to bridge these two types of omics data. For example, cTP-net(Zhou et al. 2020) is a transfer learning approach under a deep learning framework to predict surface protein levels from scRNA-seq data. The generation of diverse types of single cell data poses many computational challenges due to their high dimensionality and large sample sizes. In this paper, we focus on the annotation of single cell proteomics data to their corresponding cell types, which is a critical step in single cell analysis.

A major advantage of single cell proteomics data compared with scRNA-seq data is their high sensitivity and specificity. Based on surface marker expression patterns, manual gating can be used to identify different cell populations by the expression distributions. However, cell type labeling by manual gating on single cell data is labor intensive and subjective as it highly depends on the expert who annotates the cells. Although some methods can analyze these data based on state-of-the-art machine learning methods(Li et al. 2017; Van Gassen et al. 2015; Levine et al. 2015) for clustering cells, most of these methods are unsupervised and unable to identify cell populations automatically.

For scRNA-seq data, a number of methods have been developed to assign cells to different cell types based on expression profiles. For example, SingleR(Aran et al. 2019) trains on an extensive collection of large annotated reference transcriptomics single cell data. Preliminary labeling by SingleR can vastly accelerate cell type inference. However, there is no similar automated tool for single cell proteomics data annotation. A recently developed tool named CellGrid(Chen et al. 2020) applied the same idea to CyTOF data but needed a large number of labeled proteomics datasets as input. However, there is a lack of labeled single cell proteomics data although well-annotated scRNA-seq data are more broadly available. For the CITE-seq data, it is still necessary to annotate the transcriptomics and proteomics data separately.

To overcome the lack of labeled single cell proteomics data, we introduce an automated single cell **prot**eomics data **anno**tation approach called ProtAnno, based on non-negative matrix factorization (NMF) to incorporate data from different reference sources. The only essential input of ProtAnno is some prior knowledge on cell type-specific biomarkers. To further improve annotation accuracy, ProtAnno can take advantage of publicly available CITE-seq data and annotated scRNA-seq data. This enables ProtAnno to perform cell type annotation with no prior characterization between cell types and surface proteins by leveraging these external references.

We have evaluated the performance of ProtAnno through simulations under different settings. The results showed the robustness of ProtAnno to biological variability, technical noise, cell type number, and incomplete and inaccurate expert knowledge. We then applied ProtAnno to three real datasets: peripheral blood mononuclear cell (PBMC) paired stimulated B cell receptor CyTOF data, PBMC CITE-seq data from healthy subjects, and longitudinal whole blood covid-19 CyTOF data grouped by patients’ disease severity. In the analyses of these real data, ProtAnno provided fast and accurate labeling as demonstrated through comparisons with manual annotations or downstream biological investigations. In summary, ProtAnno is a computationally efficient and statistically robust approach for automated cell type annotation when only limited expert knowledge is available for single cell proteomics data.

## Methods

### Automated Single cell Proteomics Data Annotation Model

ProtAnno deconvolutes the proteomic expression profile *X* ∈ *R^D×C^* into the product of the cell type-specific signature matrix, *W* ∈ *R^D×K^*, and cell type assignment matrix, *H* ∈ *R^K×C^*, i.e. *X* = *WH*. In the model, we have *D* surface markers for *C* cells in the proteomics data, e.g., the cytometry data and antibody profile in CITE-seq. We denote *K* as the number of cell types. The columns of *W* are matched with the known cell types in the same order. Due to the non-negative requirement on the estimations of *W* and *H*, ProtAnno implemented NMF for solving *X* = *WH*.

In ProtAnno, we integrate information from both prior knowledge encoded in a matrix *A*_0_ and relationship between protein markers and RNA-seq data encoded in a matrix *A*. In our model, we denote *A* ∈ *R^D×G^* as the protein-RNA association matrix inferred from a CITE-seq data by elastic net [39], where *G* is the number of genes considered. A desirable CITE-seq data should include a large number of measured antibodies. ProtAnno implements two internal dictionary-like CITE-seq datasets from Unterman et al. (2020)(Unterman et al. 2020) and Ramaswamy et al. (2020) (Ramaswamy et al. 2021). Both datasets have more than 180 antibody tags. The transcriptome signature matrix *S* ∈ *R^G×K^* is generated by well-annotated scRNA-seq data. The *K* columns of *S* should be matched with *W*. We recommend using the denoised scRNA-seq data to impute drop-out events for better annotation. Specifically, we deploy SAVERx(J. Wang et al. 2019) for imputation due to its superior performance.

The expert knowledge matrix *A*_0_ ∈ *R^D×K^* is designed based on known cell type surface markers. A_0_ is a discrete matrix containing three possible values, +1, −1, and 0. If biomarker *i* should have high expression level in cell type *j*, we set 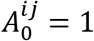; if the biomarker is not expressed in this cell type, 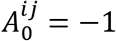; and if there is no constraint on the biomarker and cell type, then we set 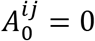.

To estimate the signature matrix *W* in the order of given cell type list, ProtAnno adds constraint on *W* with respect to the above two proteomic signature matrices, *AS* and *A*_0_.

With all the notations introduced above, the ProtAnno model is formulated as follows:

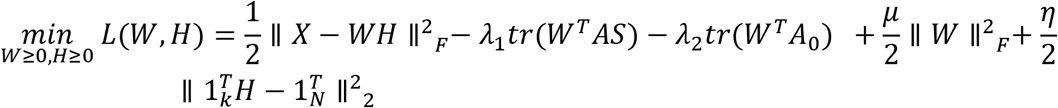

Note that ProtAnno adds regularization penalty on *W* and *H* to improve performance. We use 1_k_ and 1_N_ to denote the *K*-dimensional and *N*-dimensional column vectors with all-ones. The fourth term in ProtAnno is to control the scale of *W*, and the last term is to force the column sum of *H* to be 1.

We optimize this overall objective function based on the multiplicative update algorithm(Zhang et al. 2008; Wu and Wang 2014) to guarantee non-negativity. The algorithm requires the specifications of the penalty parameters, *λ*_l_, *λ*_2_, *μ*, and *η*. In each iteration, ProtAnno updates *W* by rows and *H* by columns. The details of ProtAnno are provided in Algorithm 1.

*Model input*: Normalized expression profile matrix *X*, discrete prior knowledge matrix *A*_0_, public labeled single cell expression profile, and penalty parameters *λ*_l_, *λ*_2_, *μ*, *η*.

*Step 1*: Set *t* = 0 and generate the initial non-negative *W* and *H* with all ones.

*Step 2*: Update *W* by rows and *H* by columns:

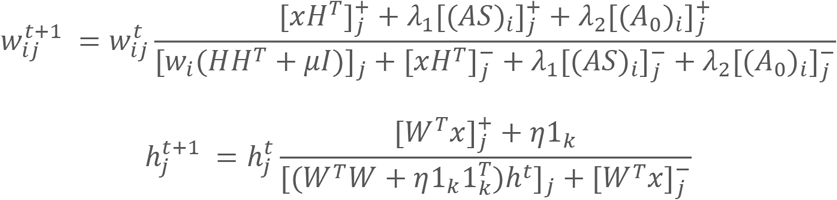

*Step 3*: Repeat Step 2 until the convergence or reaching the pre-specified number of iterations.

It can be shown that the algorithm converges in theory with the details of the theorems and converging rates provided in the supplementary materials.

### Choices of Penalty Parameters

Since the penalty parameter values are critical to ProtAnno, we developed the following algorithm to tune their values iteratively.

To ensure that ProtAnno can group expression profiles into reasonable clusters, we use the default unsupervised Louvain algorithm for preliminary clustering. We then set the initial parameter *η* by the KKT condition; and then search the initial *λ*_l_, *λ*_2_, and *μ* values by choosing from among 0.1, 1, 10, and 100. To find the initial *λ*_l_ and *λ*_2_ values, we maximize the ARI between the Louvain clusterings and ProtAnno results. We consider a novel index to select the initial value for *μ*:

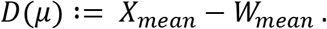

The above function considers the difference between the mean values of *X* and *W*. The main purpose of *D*(*μ*) is to ensure that *W* is on approximately the same scale as *X*.

Otherwise, the deconvolution would be unstable since we penalize the column summation of *H* to make the cell type proportion estimations meaningful.

To set more precise penalty terms, ProtAnno uses binary search to determine the final optimal output as detailed in Algorithm 2, and allows the specified search depth depending on the running time. If the final ARI is lower than 0.75, the algorithm will restart the binary search at higher resolution.

*Model input*: Normalized expression profile matrix *X*, discrete prior knowledge matrix *A*_0_, publicly labeled single cell expression profile.

*Step 1*: Estimate Louvain clusters.

*Step 2*: Initialize *W* and *H* by an optimization method of choice (*λ*_l_ = 1and *λ*_2_ = 10), and initialize *η* by KKT conditions:

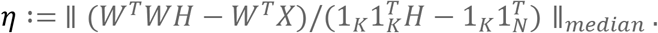

*Step 3*: Initialize *λ*_l_ and *λ*_2_ by choosing from (0.1,1,10,100) and minimize Adjusted Rank Index (ARI) with Louvain clustering.

*Step 4*: Initialize *μ* by estimated signature matrix W reliability by the following metric:

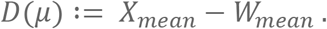

*Step 5*: Select the best *λ*_l_, *λ*_2_, and *μ* by binary search.

*Step 6* (Optional): If the final ARI with Louvain cluster is still lower than 0.75, we re-do the binary search on all the penalty parameters.

## Results

### Simulation Setup

In the following, we describe how we simulated the protein expression profiles for each single cell. We first generated the entries in the expert matrix *A*_0_ from a multinomial distribution with three categories, +1, −1, and 0, corresponding to biomarker knowledge for a cell type, i.e., high expression, no expression, and no information. The matrix will be built with the angles between vectors as large as possible, so that the simulated single cell expression profile is a nonnegative linear combination of the proteomics signature matrix(Zhang et al. 2008). To achieve it, we generate 100 matrices randomly and minmax the inner products between column vectors to get the optimal one. This design captures the nature of signature genes. However, the prior knowledge may not be in line with true relationships between markers and cell types due to technical and biological variations. We, therefore, regenerated an intermediate discrete matrix 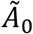 based on *A*_0_ containing three possible discrete values, 2 (high expression), 1 (low expression), and 0 (no expression), by random walk and the protocols observed in real data (details in Methods). We then derived the signature matrix *W* based on 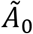 from two truncated normal distributions with high and low mean values, respectively, and varying relationships between variances and means that reflect the signal noise ratio for a biomarker. The difference of the means of the two truncated normal distributions together with their variances dictate the informativeness of a specific biomarker for cell type specification. The proteomics expression profile *X* was sampled from the truncated normal distributions based on the signature matrix that defines the mean and associated variance for each biomarker in each cell type. Finally, to simulate the predicted proteomics signature matrix from transcriptomics data, which is the product of the protein-RNA association matrix *A*, and the transcriptomics signature matrix *S*, [*AS*], we sampled the element [*AS*]_*ij*_ in the matrix with mean *W_ij_*, the expression level of signature gene *i* in cell type *j*, and variance *W_ij_*/(2 ∗ *corr*). A smaller *corr* results in a weaker correlation between the two proteomics signature matrices. All the simulation details can be found in Materials and Methods.

### Model performances and comparisons

We considered six simulation scenarios to cover a wide range of biological and technical variations in real data, with data quality varying from high (scenario 1) to low (scenario 6). We tuned the parameters of the normal distributions to decrease the cell type distinction and enlarge the expression profile divergence. To simulate the situations when AS or is too noisy to have a good prediction power, we also added more uncertainty to AS and that led to a lower correlation with real signature matrix W. The overall noise increased from scenario 1 to 6. In scenario 5, we increased the noisy level of expert matrix so that it is more aberrant compared with the single cell proteomics data expression pattern. In addition, the correlation of and was set at only 0.2. These large noises from references would reduce the annotation accuracy. Furthermore, we increased expression profile variations by lowering the mean-variance ratio in scenario 6, making the annotation more challenging. All the parameters settings and details are listed in Supplementary Table 1.

We compared the performance of six models: the full ProtAnno model, the ProtAnno model without transcriptomics information *AS*, the ProtAnno model without expert knowledge *A*_0_, the unsupervised ProtAnno model, the unsupervised Louvain clustering, and cell type assignment by non-negative least squares (NNLS) when the protein signature matrix *W* is known. The last one is the best result an NMF model can achieve. We consider five metrics for annotation accuracy: Adjusted Rand Index (ARI), annotation accuracy assigned by the largest value for every single cell, Normalized Mutual Information (NMI), cosine similarity between the estimated and real assignment, and Average Silhouette Width.

The simulation results in Figure 2A show that the full ProtAnno model achieved the best annotation accuracy, especially compared with the NNLS model. Specifically, from scenarios 1 to 4, ProtAnno was able to achieve the same performance an NNLS model could have when the true signature matrix is known. These results suggest the importance of having precise information for *A*_0_ and *AS*. Besides the full ProtAnno model, the partial ProtAnno model with the expert matrix had the second-best performance, just slightly worse than the full model. The partial ProtAnno model only having the transcriptomics reference was not competitive with the first two models. But it was still much better than the unsupervised clustering, even in terms of ARI and NMI. In general, both full and partial ProtAnno models could obtain accurate cell type assignments even in the presence of considerable noise.

**Fig1.**
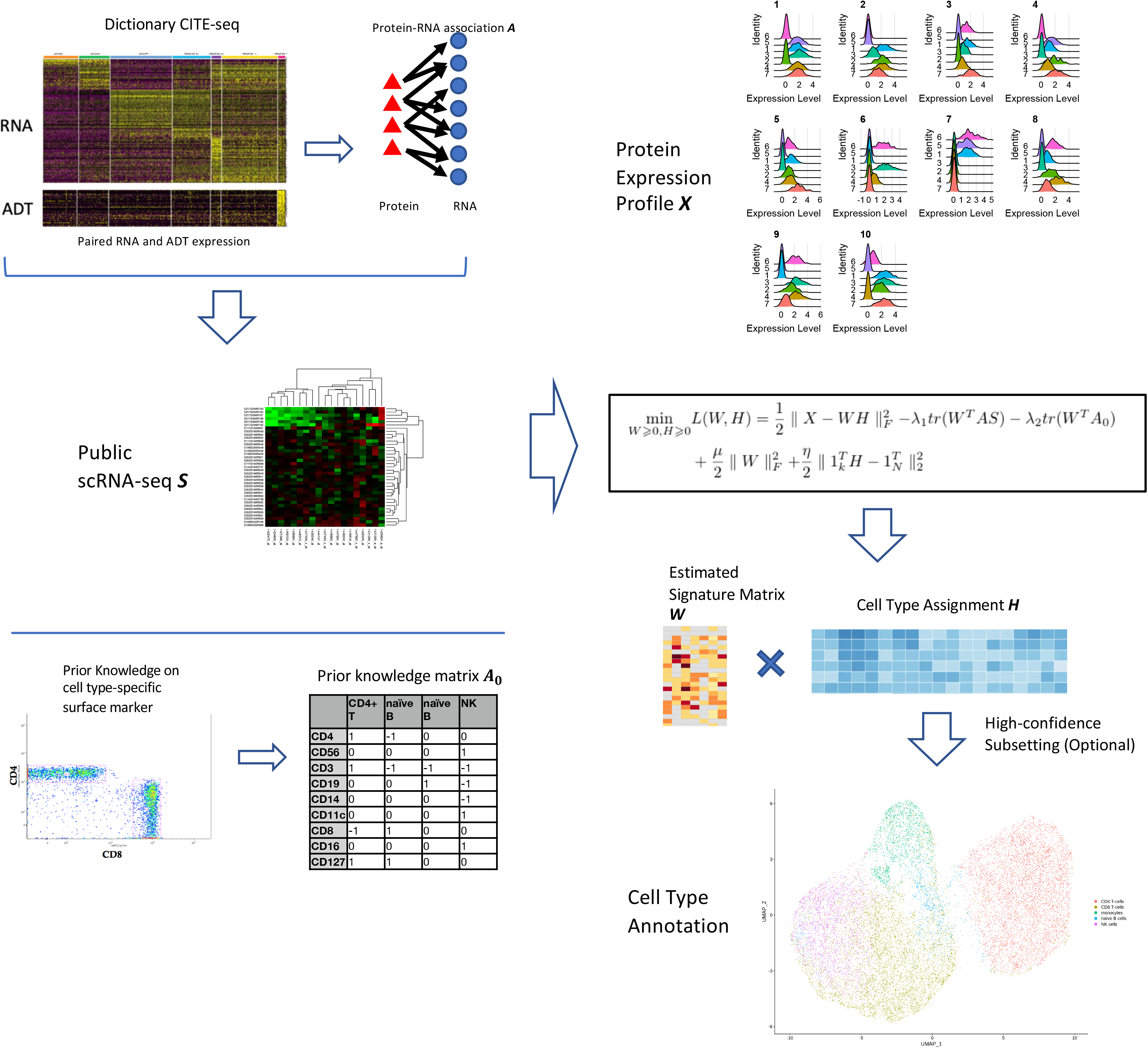
Model overview. ProtAnno first generates a protein-RNA association matrix A trained on CITE-seq dictionary by Elastic Net and a transcriptome signature matrix *S* from scRNA-seq with cell type annotations. The product of *A* and *S* is the predicted protein level signature matrix based on mRNA counts. The other input is the discrete matrix *A*_0_ to represent the general expert biomarker knowledge. The loss function is an NMF optimization program on protein expression matrix to deconvolute into estimated protein signature matrix *W* and cell type assignment matrix *H*, with penalizations on *AS* and *A*_0_ and regularizations on *W* scale and column sum of *H*. The matrix *H* could be optionally trimmed to the cells which have high-confidence assignment in the presence of high noise in *X*. The final annotations of ProtAnno are based on the largest value of *H* column-wise.

**Fig2.**
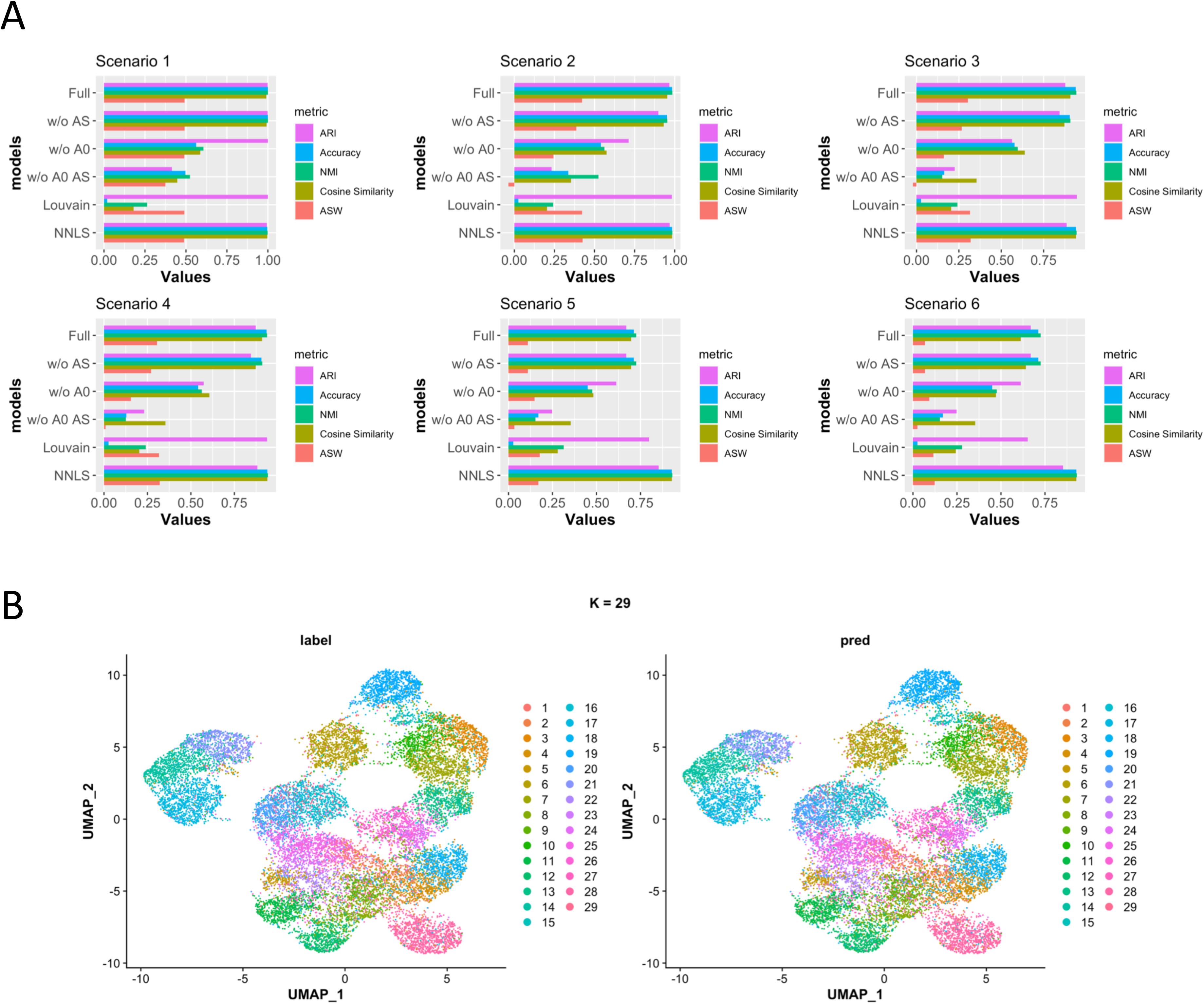
ProtAnno simulation benchmarking results. A) Comparisons of benchmarking annotation results under six simulation models. The x-axis lists the six models; the y-axis shows the annotation metric values. B) An example of ProtAnno annotation when the number of cell types *K* = 29. The left panel is the UMAP plot on true labels; the right panel is the UMAP plot of ProtAnno annotations.

We investigated the numerical convergence of ProtAnno in our simulation studies. Fig 2B shows an example of ProtAnno cell type assignment in a UMAP plot with 29 cell types, where the accuracy was 98.8%. The high agreement of clustering between the real labels (left) and the ProtAnno annotation (right) demonstrates the power of ProtAnno.

### Algorithm convergence and robustness

ProtAnno generally converges after 100 iterations (Supp 1A). To filter the high confidence annotation, the subsetting step will keep the cell whose estimated *H* is greater than 0.5 after column normalization. The subsetting step was able to slightly enhance the annotation accuracy (Supp 1B).

ProtAnno’s robustness was investigated with respect to three parameters: mean-variance ratio (Fig 3A), correlation between *W* and *AS* (Fig 3A, supp 1C), and cell type number *K* (Fig 3C). Since a protein with a higher mean expression level tends to have a higher variance, we used mean-variance ratio to characterize the signal noise ratio of a protein. As expected, the prediction accuracy dropped with a reduced mean-variance ratio (Fig 3A). Overall, the full ProtAnno model was very robust even when the *AS* prediction power was weak (supp 1C), implying the critical role of *A*_0_. We also explored how the transcriptomics information AS could help the annotation when *A*_0_ is not available (supp 1C). The accuracy rate was almost linearly associated with the prediction correlation. Therefore, scRNA-seq data and dictionary CITE-seq data with high quality are essential for the good performance of the partial ProtAnno model when no reliable expert knowledge is available. As for the impact of the number of cell types, ProtAnno was robust to the large cell type number as long as the cell counts of each cell population were more than 100 in our simulation (Fig 2B, Fig 3C).

**Fig3.**
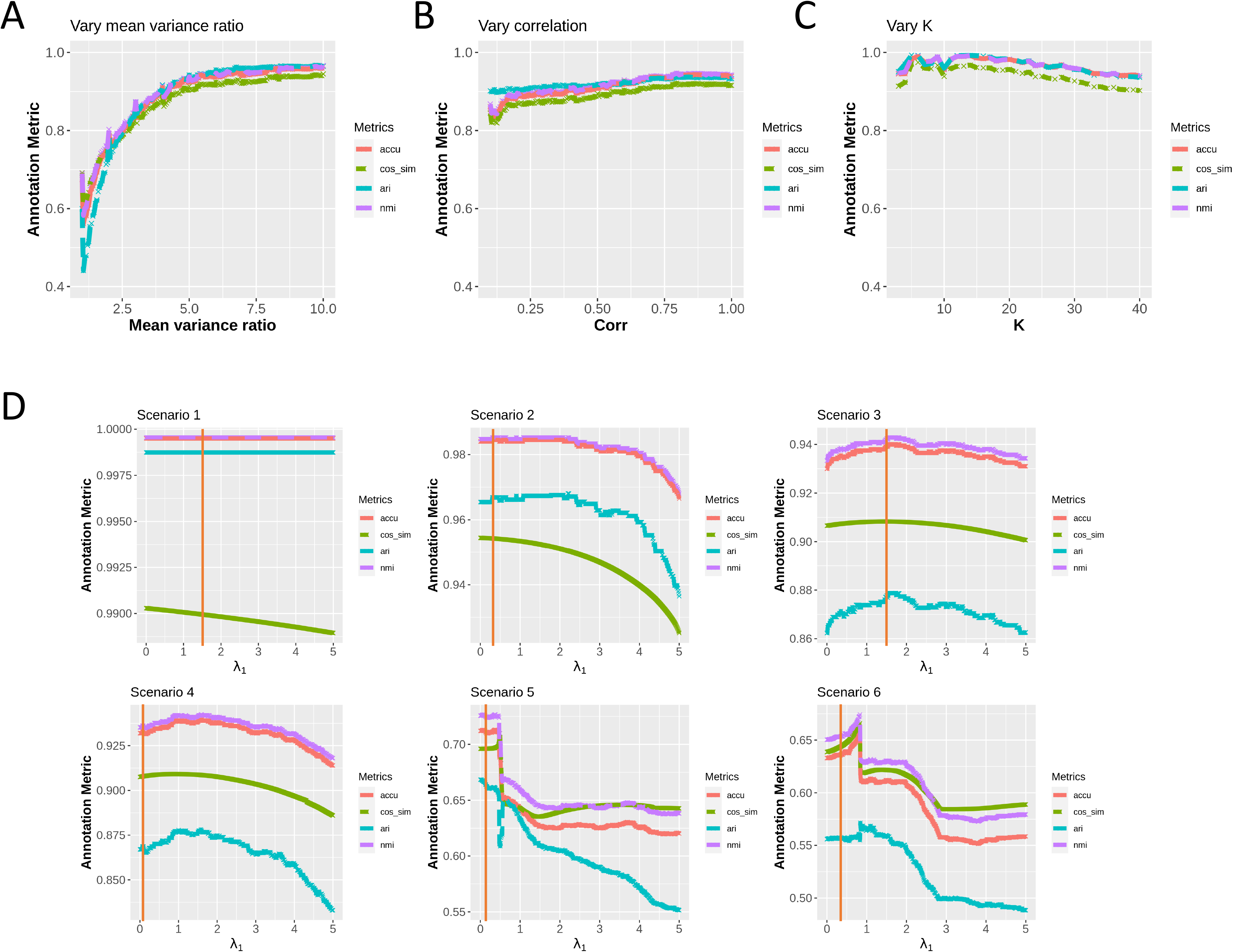
ProtAnno model robustness evaluations. A) The relationship between ProtAnno annotation accuracy and expression variation. The x-axis represents the expected ratio of mean and variance. A higher value indicates lower expression variation. The y-axis shows the values of annotation metrics. The curves represent the results from 100 simulations for each mean-variance ratio value. B) The effect of transcriptomics data on a full ProtAnno model. The x-axis represents the correlation between transcriptomics predicted signature matrix *AS* and real protein signature matrix *W*. The y-axis represents the value of annotation metrics. C) ProtAnno robustness as a function of cell type number *K*. The x-axis represents cell type number that varies from 3 to 40. The y-axis represents the value of annotation metrics. D) Evaluation of parameter tuning algorithm. The x-axis represents the change of *λ*_l_ from 0 to 5 under six simulation scenarios. The color represents different annotation metrics. The y-axis represents the value of annotation metrics. The red vertical line represents the optimal *λ*_l_ by the parameter tuning algorithm for each simulation scenario.

The good performance of ProtAnno depends on the appropriate choices of the penalty parameters. Simulation results suggest that our proposed parameter tuning algorithm was able to find good penalty values (Fig 3D, Supp 1D) as shown in the relationship between algorithm performance and tuning parameter values in these figures in most cases.

Finally, we can rewrite the ProtAnno objective function in an equivalent form as (see supplementary materials):

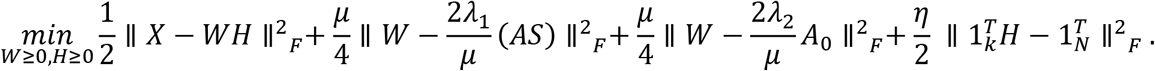

The above objective function form illustrates the imposed relationships between W, AS, and *A*_0_. The second penalty terms will force the ratio of elements in matrices W and AS to be approximately equal to 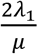. Specifically, the ratio of *W_ij_*/[*AS*]_*ij*_ should be approximately 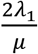 for any entry with row index *i* and column index *j*. Similarly, the ratio of *W* and *A*_0_ should be approximately 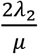. Therefore, we empirically examined whether the ratios of matched elements in matrices W and AS (or *A*_0_) approached the theoretical ones (Supp 2A, 2B). The figures show that the empirical distribution of these ratios compared with 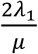 and 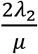, suggesting that estimated *W* was close to 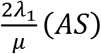 and 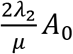 at the same time.

### Application to paired stimulated B cell receptor/Fc receptor cross-linker (BCR) cytometry data

We first evaluated ProtAnno’s performance on a benchmarking BCR CyTOF dataset with eight paired samples from PBMC in Bodenmiller et al. (2012)(Bodenmiller et al. 2012). Compared with the reference group, the B cell population was stimulated by BCR in the treatment group. All the samples were collected from healthy individuals, and 10 cell type markers were measured in a total of 172,791 cells. This dataset was labeled in Nowicka et al. (2017)(Nowicka et al. 2017) after the authors applied an unsupervised clustering method FlowSOM(Van Gassen et al. 2015) and manually merged clusters. We evaluated the assignments of these cells to five major cell types based on the 10 cell type markers. Other inputs, the association matrix and the transcriptomics signature matrix S, were generated by the CITE-seq data in Ramaswamy et al. (2020)(Ramaswamy et al. 2021). To match with annotated cell types by Nowicka et al., we manually gated the five major cell types in the scRNA-seq. We also constructed an expert-guided matrix *A*_0_ (Table 1). ProtAnno assigned a cell to the cell type with the largest value in matrix H.

**Table 1.**

| Marker | memory B cells | naïve B cells | CD4 T-cells | CD8 T-cells | DC | monocytes | NK cells |
| --- | --- | --- | --- | --- | --- | --- | --- |
| CD3 | -1 | -1 | 1 | 1 | -1 | -1 | -1 |
| CD45 | 1 | 1 | 1 | 1 | 1 | 1 | 1 |
| CD4 | 0 | 0 | 1 | -1 | 0 | 0 | 0 |
| CD20 | 1 | 1 | 0 | 0 | -1 | -1 | -1 |
| CD33 | 0 | 0 | 0 | 0 | 0.5 | 1 | 0 |
| CD123 | 0 | 0 | 0 | 0 | 1 | 0 | 0 |
| CD14 | 0 | 0 | 0 | 0 | -1 | 1 | -1 |
| IgM | 1 | -1 | 0 | 0 | 0 | 0 | 0 |
| HLA-DR | 0 | 0 | 0 | 0 | 1 | 0 | -1 |
| CD7 | 0 | 0 | 1 | 1 | 0 | 0 | 1 |

The overall median accuracy of ProtAnno for this data set was 83% (Fig 4A) across the samples. For most of the samples, the accuracy could be as high as 80% to 85% (Supp3), except an outlier, patient 8, with decreased the average. For this data set, even the partial ProtAnno with only *A*_0_ or only *AS* could achieve an accuracy of over 70% overall. The subsetting step could slightly improve accuracy by keeping high-confidence cells (Supp 3, Supp 4). Because the BCR dataset does not have clear clustering patterns, making it challenging to annotate cells. This can be seen in the UMAP plot, where some of the cells from different cell types are located in the same region (Fig 4E). Similar observation was made in the benchmarking study of unsupervised clustering accuracy by Weber et al. (2019)(Weber et al. 2019). At the lowest clustering resolution of 9 clusters, the false positive rate was around 20%, where unsupervised clustering could not achieve accurate annotation. Therefore, an overall accuracy of 80% may be considered satisfactory for this dataset. We further investigated the misclassification patterns by the confusion matrix for patient 1 in the stimulated group (Fig 4B). It can be seen that most of the cells could be correctly assigned by ProtAnno. However, some of the natural killer cells (NK cells) were annotated as CD8 T cells. That is partly because the BCR data did not measure the biomarker CD8, which offers important information for CD8 T cell assignment. For this data set, the only biomarker that was informative to distinguish these two cell populations is CD3, which makes computational annotation difficult. The UMAP plot also shows that the annotation is highly consistent with the manually annotated cell types, although the boundary of NK cells and CD8 T cells is hard to recognize (Fig 4E).

**Fig4.**
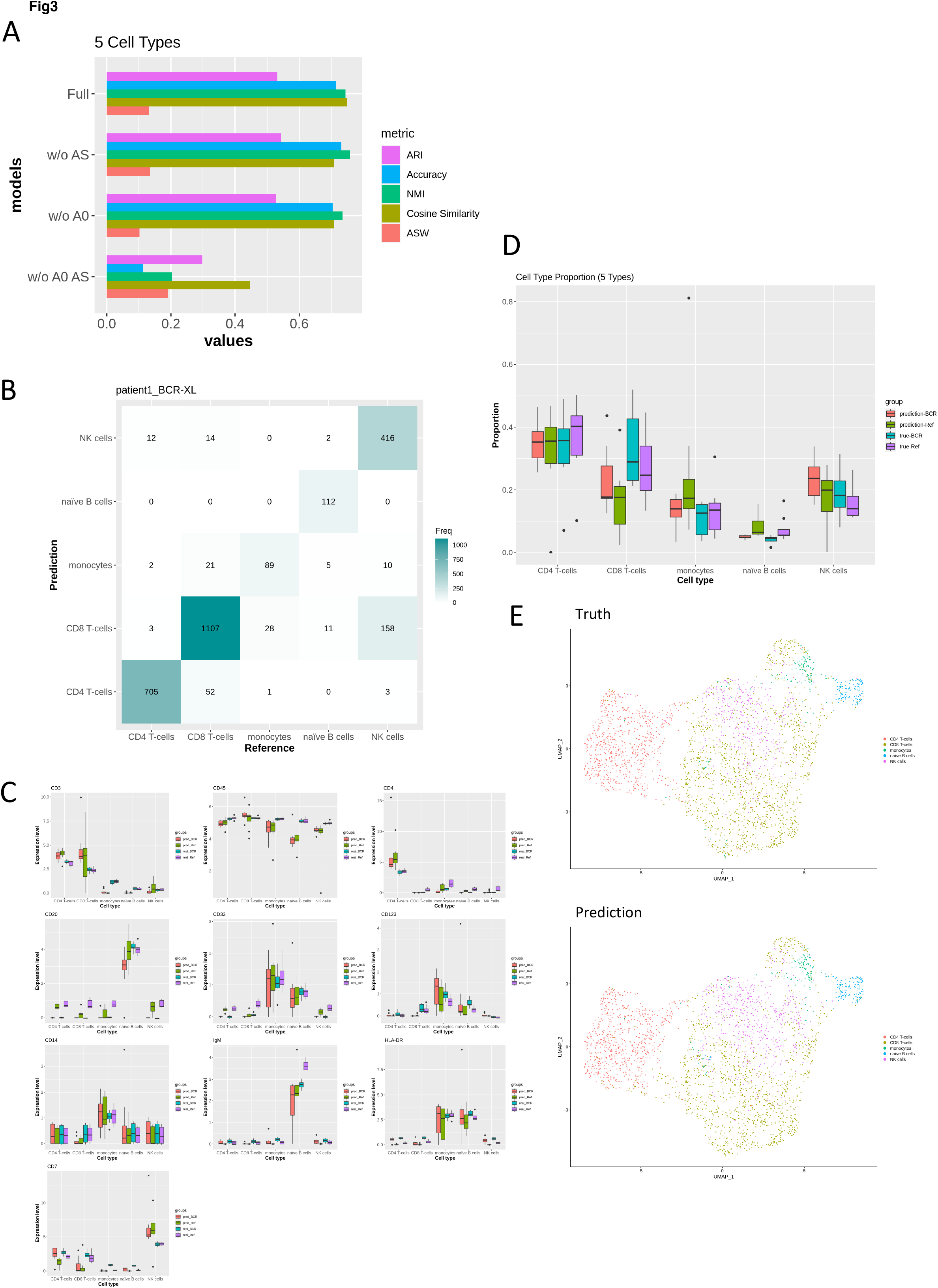
ProtAnno results on BCR cytometry data. A) Benchmarking ProtAnno on five labeled cell types. The x-axis lists the four ProtAnno models; the y-axis is the annotation metric value. B) Confusion matrix of ProtAnno annotation on BCR stimulated patient 1. The y-axis is the ProtAnno results compared with the true cell type labels as a reference in the x-axis. C) The boxplots of estimated signature expression in *W*. The x-axis represents the cell type and the y-axis is the distribution of values in the signature matrix across patients within groups. The first two columns are ProtAnno annotation results in the stimulated and unstimulated BCR groups and the last two columns are the true average signature expression distributions. D) Cell type proportions boxplots. In this figure, we compare cell type proportions estimated by ProtAnno with true proportions. The first two columns are ProtAnno annotation results in the stimulated and unstimulated BCR groups and the last two columns are the true cell type proportions in these two groups. E) The UMAP plot. The upper panel is the true annotation; the lower panel is the assignment by ProtAnno.

Since many genes had significant differential expression levels across groups in the BCR dataset, we studied the W matrix variations to test whether ProtAnno could accurately estimate the signature matrix. The most differential expression directions in the stimulated group by ProtAnno are consistent with prior biological knowledge (Fig 4C). For example, biomarker HLA-DR had a significantly elevated expression in most cell types after BCR stimulation. The inferred cell type proportions also suggest that ProtAnno can provide an accurate estimation in the comparative study. For example, the proportions of naive B cells were lower in the stimulated group (Fig 4D). These downstream analyses suggest accurate annotation by ProtAnno.

### Application to PBMC CITE-seq data

To investigate the performance of ProtAnno on the proteomic expression profile in CITE-seq data, we analyzed a PBMC CITE-seq dataset reported in Wang et al. (2020)(X. Wang et al. 2020). This dataset measured 10 surface markers in 1372 cells from a healthy subject. We used ProtAnno to annotate 1153 cells with six cell types, because we cannot annotate the other 219 cells to any cell type based on the available limited 10 markers. These cells were manually labeled and the details are provided in the supplementary materials. We used the CITE-seq data in Unterman et al. (2020)(Unterman et al. 2020) to derive the transcriptomics signature matrix and the CITE-seq data in Ramaswamy et al. (2020)(Ramaswamy et al. 2021) to obtain the association matrix and construct *A*_0_(Table 2). The automated annotation accuracy was around 99% with 100% accuracy for most cell types, except for non-classical monocytes. ProtAnno failed to recognize this cell type since it is a rare population whose cell type proportion is below 1% (Fig 5A). These results suggest that ProtAnno could be applied to multiple single cell proteomics sequencing platforms, though cytometry and CITE-seq are different technologies with distinct features.

**Table 2.**

| Marker | CD4+T cell | CD8+T cells | naïve B cells | NK cell | classical monocytes | non-classical monocytes |
| --- | --- | --- | --- | --- | --- | --- |
| CD154 | 0 | 0 | 0 | 0 | 1 | 1 |
| CD4 | 1 | -1 | 0 | 0 | 0 | 0 |
| CD56 | 0 | 0 | 0 | 1 | -1 | -1 |
| CD3 | 1 | 1 | -1 | -1 | -1 | -1 |
| CD19 | 0 | 0 | 1 | -1 | -1 | -1 |
| CD14 | 0 | 0 | 0 | -1 | 1 | -1 |
| CD11c | 0 | 0 | 0 | 1 | 0 | 0 |
| CD8 | -1 | 1 | 0 | 0 | 0 | 0 |
| CD16 | 0 | 0 | 0 | 1 | -1 | 1 |
| CD127 | 1 | 1 | 0 | 0 | 0 | 0 |

**Fig5.**
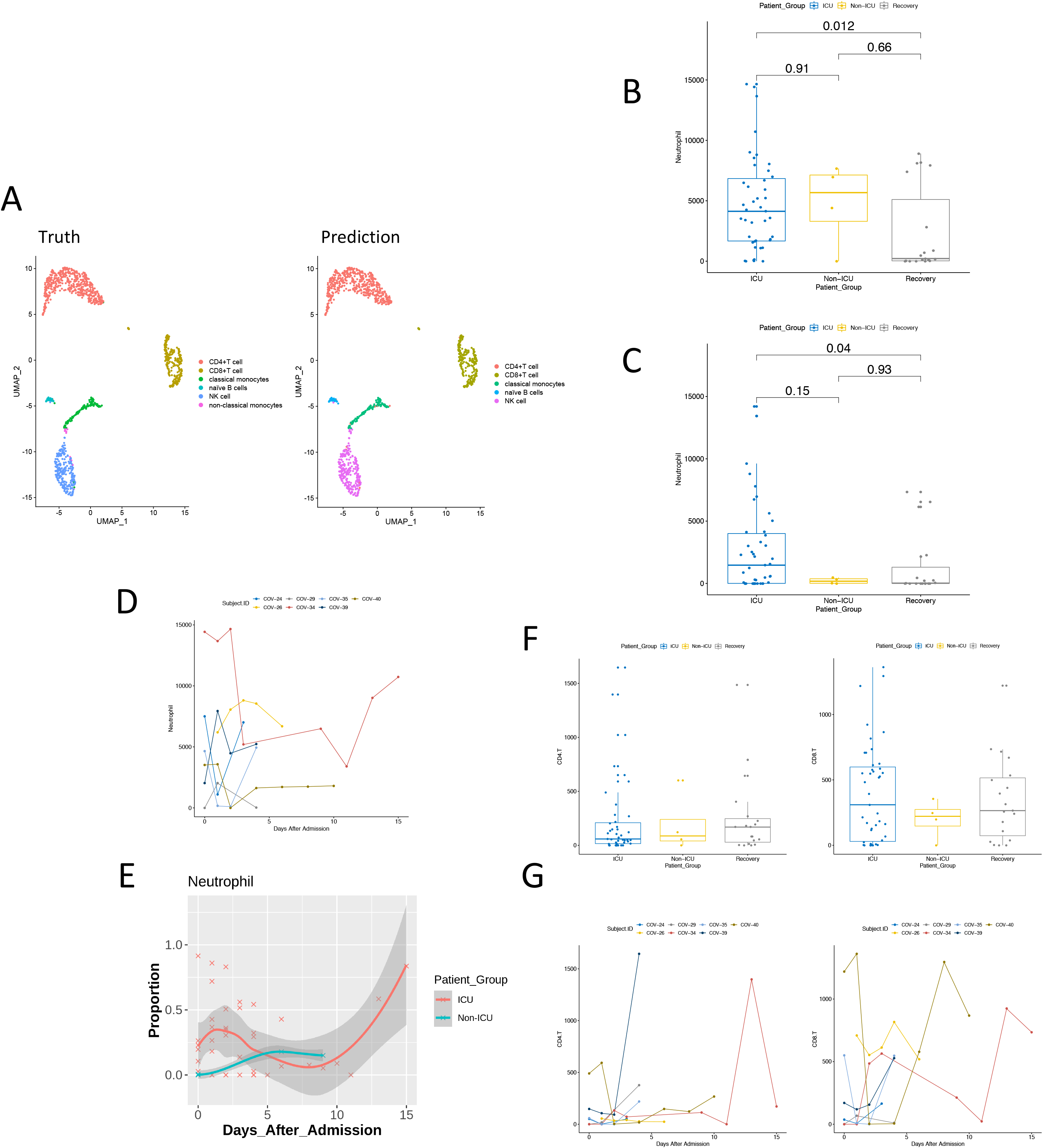
ProtAnno results on PBMC CITE-seq data and temporal whole blood covid-19 cytometry data. A) The UMAP plot of the proteomic data in the PBMC CITE-seq study in Wang et al. (2020)(X. Wang et al. 2020). The left panel is the true annotation; the right panel is the assignment by ProtAnno. The UMAP projections are colored by the cell types. B) and C) Boxplots of cell type counts of neutrophil cells from the whole blood covid-19 study in Rodriguez et al. (2020)(Rodriguez et al. 2020) by ProtAnno across different patient groups i.e., ICU, non-ICU, and recovery patients. B) is the raw output of ProtAnno. C) is the subsetted output of ProtAnno for keeping high-confidence assignments only. The paired Wilcoxon test p-values are shown on the top of the figures. D) Neutrophil cell counts estimation by ProtAnno over the days after admission. The color represents the patient who has the longitudinal cytometry data. E) Neutrophil cell type proportions estimated by ProtAnno over the days after admission. Two curves represent the ICU and non-ICU patient groups. F) Boxplots of CD4 T cell and CD8 T cell counts across groups (ICU, non-ICU, and recovery patients). G) CD4 T and CD8 T cell type proportions estimation by ProtAnno over the days after admission. The color represents the patient who has the longitudinal cytometry data.

### Application to longitudinal whole blood covid-19 cytometry data

Finally, we applied ProtAnno to a longitudinal whole blood covid-19 CyTOF data set reported in Rodriguez et al. (2020)(Rodriguez et al. 2020). This dataset included 37 adult covid-19 patients classified into ICU, non-ICU, and recovery groups. All the hospitalized (ICU and non-ICU) patients were measured at multiple time points. Each sample had 15,000~20,000 cells. The authors used an in-house annotation method CellGrid(Chen et al. 2020) to train a subtyping strategy with high resolution to annotate over 50 cell types. Since we do not have access to such a large training CyTOF dataset and comprehensive knowledge on biomarkers, we selected 23 surface markers to define 17 common cell types in whole blood. For this dataset, we only considered the partial ProtAnno model with *A*_0_ since there is no whole blood CITE-seq dataset available (Table 3). Since we do not have ground truth on cell type annotations, we primarily evaluated the performance of ProtAnno through downstream analysis. More specifically, we studied the cell counts and cell types of neutrophils and lymphocytes.

**Table 3.**

| marker | CD8 T | CD4 T | Mem Treg | Naive Treg | gdT | Naive B | Memory B | Plasmablast | NK | pDC | mDC | Monocyte | Eosinophil | Neutrophil | MAIT | CD16+ Basophil | CD16- Basophil |
| --- | --- | --- | --- | --- | --- | --- | --- | --- | --- | --- | --- | --- | --- | --- | --- | --- | --- |
| CD3 | 1 | 1 | 1 | 1 | 1 | -1 | -1 | -1 | -1 | -1 | -1 | -1 | -1 | -1 | 1 | -1 | -1 |
| IgD | 0 | 0 | 0 | 0 | 0 | 1 | -1 | 0 | 0 | 0 | 0 | 0 | 0 | 0 | 0 | 0 | 0 |
| CD20 | 0 | 0 | 0 | 0 | 0 | 1 | 1 | -1 | -1 | -1 | -1 | -1 | 0 | 0 | 0 | 0 | 0 |
| CD4 | -1 | 1 | 1 | 1 | -1 | 0 | 0 | 0 | 0 | 0 | 0 | 0 | 0 | 0 | -1 | 0 | 0 |
| CD8a | 1 | -1 | -1 | -1 | -1 | 0 | 0 | 0 | 0 | 0 | 0 | 0 | 0 | 0 | 0 | 0 | 0 |
| CD11c | 0 | 0 | 0 | 0 | 0 | 0 | 0 | 0 | 0 | 1 | -1 | 0 | 0 | 0 | 0 | 0 | 0 |
| CD16 | 0 | 0 | 0 | 0 | 0 | 0 | 0 | 0 | 0 | -1 | -1 | 0 | 0 | 1 | 0 | 1 | -1 |
| CD14 | 0 | 0 | 0 | 0 | 0 | 0 | 0 | 0 | -1 | -1 | -1 | 1 | 0 | -1 | 0 | 0 | 0 |
| CD45RA | 0 | 0 | -1 | 1 | 0 | 0 | 0 | 0 | 0 | 0 | 0 | 0 | 0 | 0 | 0 | 0 | 0 |
| CD127 | 1 | 1 | -1 | -1 | 0 | 0 | 0 | 0 | 0 | 0 | 0 | 0 | 0 | 0 | 0 | 0 | 0 |
| CD33 | 0 | 0 | 0 | 0 | 0 | 0 | 0 | 0 | 0 | 0 | 0 | 1 | 0 | 0 | 0 | 1 | 1 |
| CD24 | 0 | 0 | 0 | 0 | 0 | 1 | 1 | 0 | 0 | 0 | 0 | 0 | 1 | 0 | 0 | 0 | 1 |
| TCRgd | 0 | 0 | 0 | 0 | 1 | 0 | 0 | 0 | 0 | 0 | 0 | 0 | 0 | 0 | 0 | 0 | 0 |
| CD123 | 0 | 0 | 0 | 0 | 0 | 0 | 0 | 0 | 0 | -1 | 1 | 0 | 0 | 0 | 0 | 1 | 1 |
| CD56 | 0 | 0 | 0 | 0 | 0 | 0 | 0 | 0 | 1 | -1 | -1 | -1 | -1 | -1 | 1 | 0 | 0 |
| CD25 | -1 | -1 | 1 | 1 | 0 | 0 | 0 | 0 | 0 | 0 | 0 | 0 | 0 | 0 | 0 | 0 | 0 |
| HLA-DR | 0 | 0 | 0 | 0 | 0 | 0 | 0 | 0 | -1 | 1 | 1 | 0 | 0 | 0 | 0 | 0 | 0 |
| CD161 | 0 | 0 | 0 | 0 | 0 | 0 | 0 | 0 | 0 | 0 | 0 | 0 | 0 | 0 | 1 | 0 | 0 |
| CD28 | 0 | 0 | 1 | 1 | 0 | 0 | 0 | 0 | 0 | 0 | 0 | 0 | 0 | 0 | 0 | 0 | 0 |
| CD38 | 0 | 0 | 0 | 0 | 0 | 0 | 0 | 1 | 1 | 0 | 0 | 0 | 0 | 0 | 0 | 0 | 0 |
| CD27 | 0 | 0 | 0 | 0 | 0 | -1 | 1 | 0 | 0 | 0 | 0 | 0 | 0 | 0 | 0 | 0 | 0 |
| CD55 | 0 | 0 | 0 | 0 | 0 | 0 | 0 | 0 | 0 | 0 | 0 | 0 | 0 | 1 | 0 | 0 | 0 |
| Siglec-8 | 0 | 0 | 0 | 0 | 0 | 0 | 0 | 0 | 0 | 0 | 0 | 0 | 1 | 0 | 0 | 1 | 1 |

The neutrophiles are strongly positively correlated with patient disease severity (Aschenbrenner et al. 2021). In this dataset, we treated the patient group as the severity indicator. In the ProtAnno results, the neutrophil cell counts of recovery patients were significantly lower than those of the hospitalized patients (Fig 5B, 5C) with the Wilcoxon test p-value smaller than 0.05 for both raw output and subsetting output of ProtAnno, consistent with our expectation. It is also known that the inflammatory symptoms are dramatically elevated in severe cases(C. Huang et al. 2020; Ong et al. 2020; Su et al. 2020; Hadjadj et al. 2020). Longitudinal analysis of this dataset showed a slight decrease of neutrophils proportions over time after admission, except with one patient, COV-34, having a slight increase at the end of the study (Fig 5D, 5E). The changes between day 1 to day 12 were significant, especially for the ICU group with a p-value of 0.029. These results suggest that most ICU patients were recovered after treatment.

In contrast, the lymphocyte proportion is expected to decline in severe covid-19 patients. Lymphopenia has been used as a biomarker to define a patient's morbidity(Fathi and Rezaei 2020; Tavakolpour et al. 2020; I. Huang and Pranata 2020). When we applied ProtAnno to infer CD4 T and CD8 T cells, although there was no statistically significant difference across groups, the CD4 T cell count was indeed elevated if the patients recovered (Fig 5F). In the longitudinal curves of cell counts, almost all the patients had increased CD4 T cell and CD8 T cell counts after treatment (Fig 5G). This implies that the lymphopenia was alleviated by treatment.

The analysis of this data set shows the usefulness of ProtAnno even when the data are highly noisy.

## Discussion

In this paper, we have developed a computationally efficient and statistically robust method, ProtAnno, for automated annotation for single cell proteomic data. Using protein expression profiles, this computationally efficient method annotates cells based on cell type information gathered through (imprecise) prior knowledge, publicly annotated single cell datasets, and CITE-seq data. ProtAnno only requires simple and accessible references, e.g. publicly annotated scRNA-seq. When only limited references are available, ProtAnno can be applied either without prior biological knowledge or public transcriptomics references. ProtAnno can also resolve heterogeneity among cell populations. For instance, ProtAnno was able to detect signature matrix variations across different sample groups in the BCR study. In real data applications, we showed that we can gain biological insights through downstream analysis, such as cell type-specific differential expression analysis and cell type proportion comparisons after annotations by ProtAnno.

Simulation results showed that ProtAnno is robust to the number of cell types and noises in cytometry and CITE-seq antibody expression profiles. However, ProtAnno may be limited to major cell types and unstable when the data are very noisy. Specifically, it is common that the single cell data does not have a clear separated clustering. For example, ProtAnno does not always have a reliable assignment for all samples (supp 3). When a specific cell type is hard to recognize even for manual gating, ProtAnno may misclassify its entire cell subpopulation. The annotation performance also dropped substantially in simulations with large noise in X and/or more limited information from references *A*_0_ and *AS*. Therefore, ProtAnno may be suitable for preliminary assignments and may require manual inspection.

Future work can incorporate hierarchical cell type structure to enhance the classification resolution and accuracy. Specifically, ProtAnno could be extended with multiple layers for classification. The cell populations with low resolution, i.e., neutrophils and lymphocytes, could be identified first, and the cell subtypes, i.e., naive CD4 T cell and memory CD8 T cell, can be classified in the next few steps. Additionally, some steps in the parameter tuning algorithm could be replaced by advanced computational clustering methods, i.e., FlowSOM(Van Gassen et al. 2015) and SPADE(Qiu et al. 2011). This may be helpful when the boundaries of cell types are unclear. Lastly, although ProtAnno does not need training data, a more unbiased association matrix inferred from more data could be a helpful add-on item to ProtAnno.

In summary, ProtAnno can provide a robust preliminary automated cell type annotation. Its performance could be further improved through adopting more advanced clustering approaches and more reliable references.

**Table 4.**
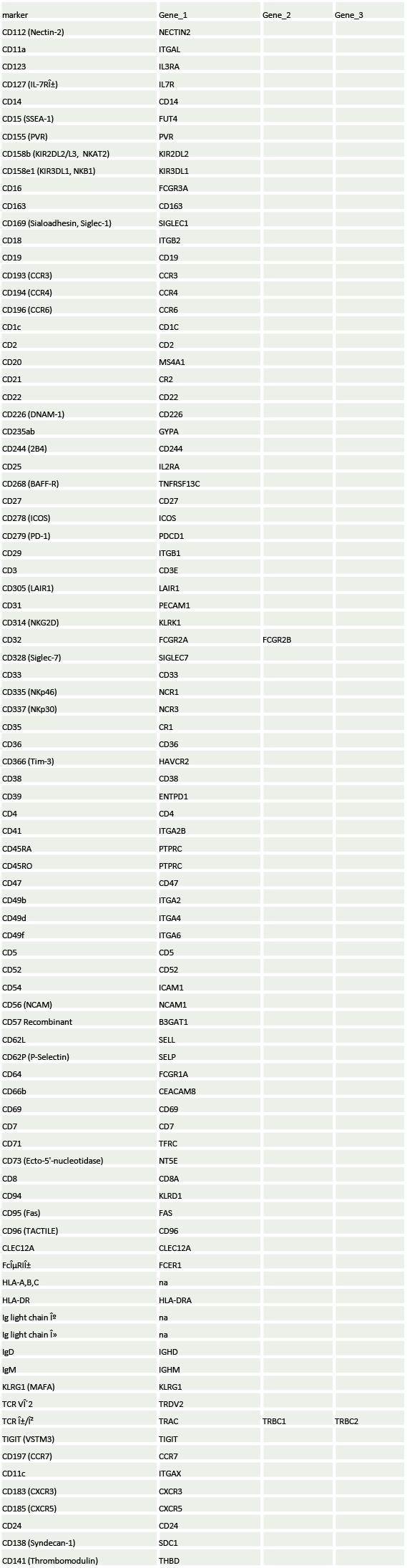

## Figures and Legends

**Supp1.**
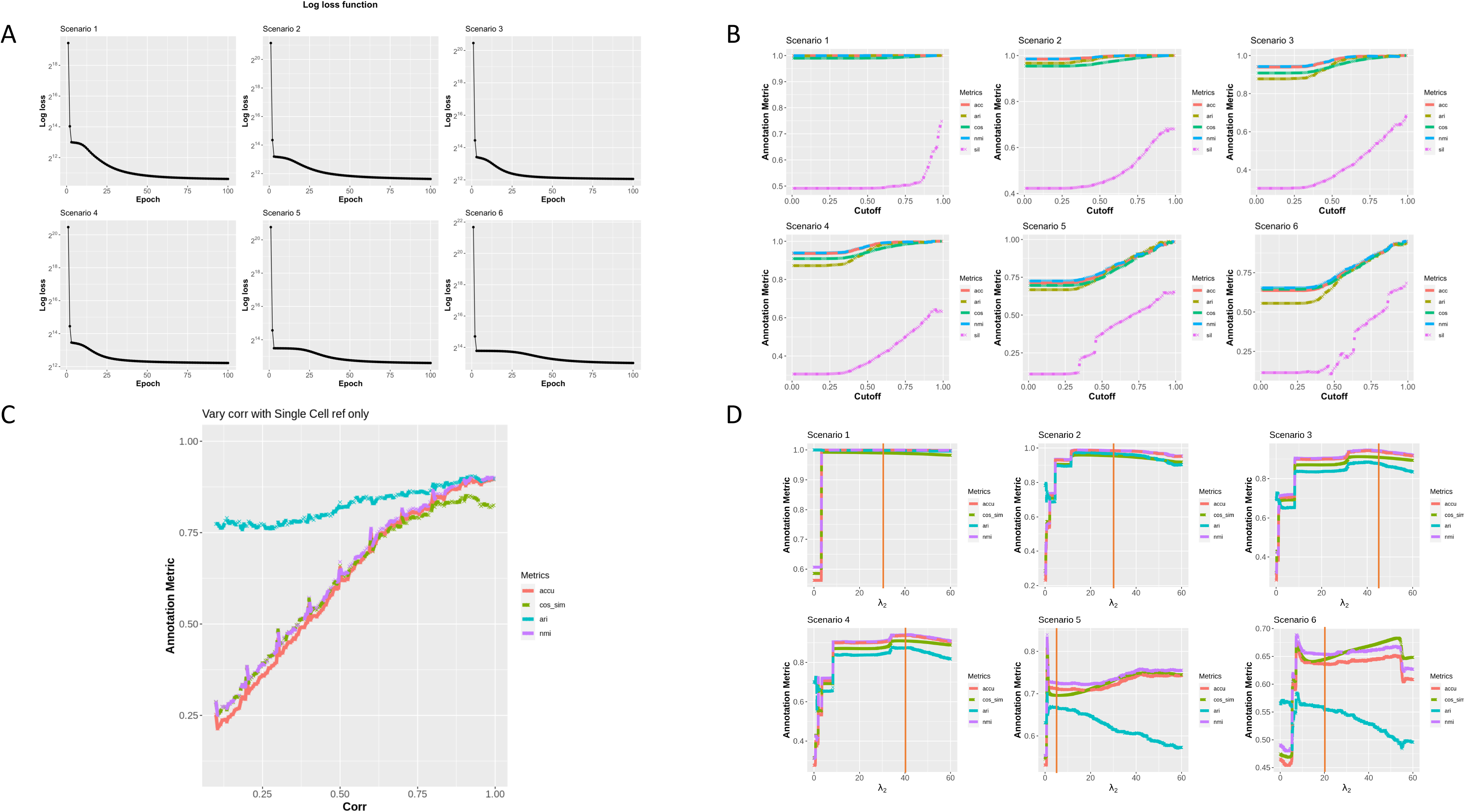
ProtAnno optimization properties. A) ProtAnno convergence for six simulation scenarios. The x-axis represents iteration number. The y-axis represents log loss function values. B) The subsetting cutoff parameter. The x-axis represents the cutoff value. The y-axis represents annotation metrics value. C) The impact of transcriptomics data on annotation in the ProtAnno model with only transcriptomics penalization, *AS*. The x-axis represents the correlation between transcriptomics predicted signature matrix *AS* and true protein signature matrix *W*. The y-axis represents the annotation metrics value. D) The evaluation of parameter tuning algorithm. The x-axis corresponds to *λ*_2_ from 0 to 60 under six simulation scenarios. The color represents different annotation metrics. The y-axis represents the annotation metrics value. The red vertical line represents the optimal *λ*_2_ by the parameter tuning algorithm for each simulation scenario.

**Supp2.**
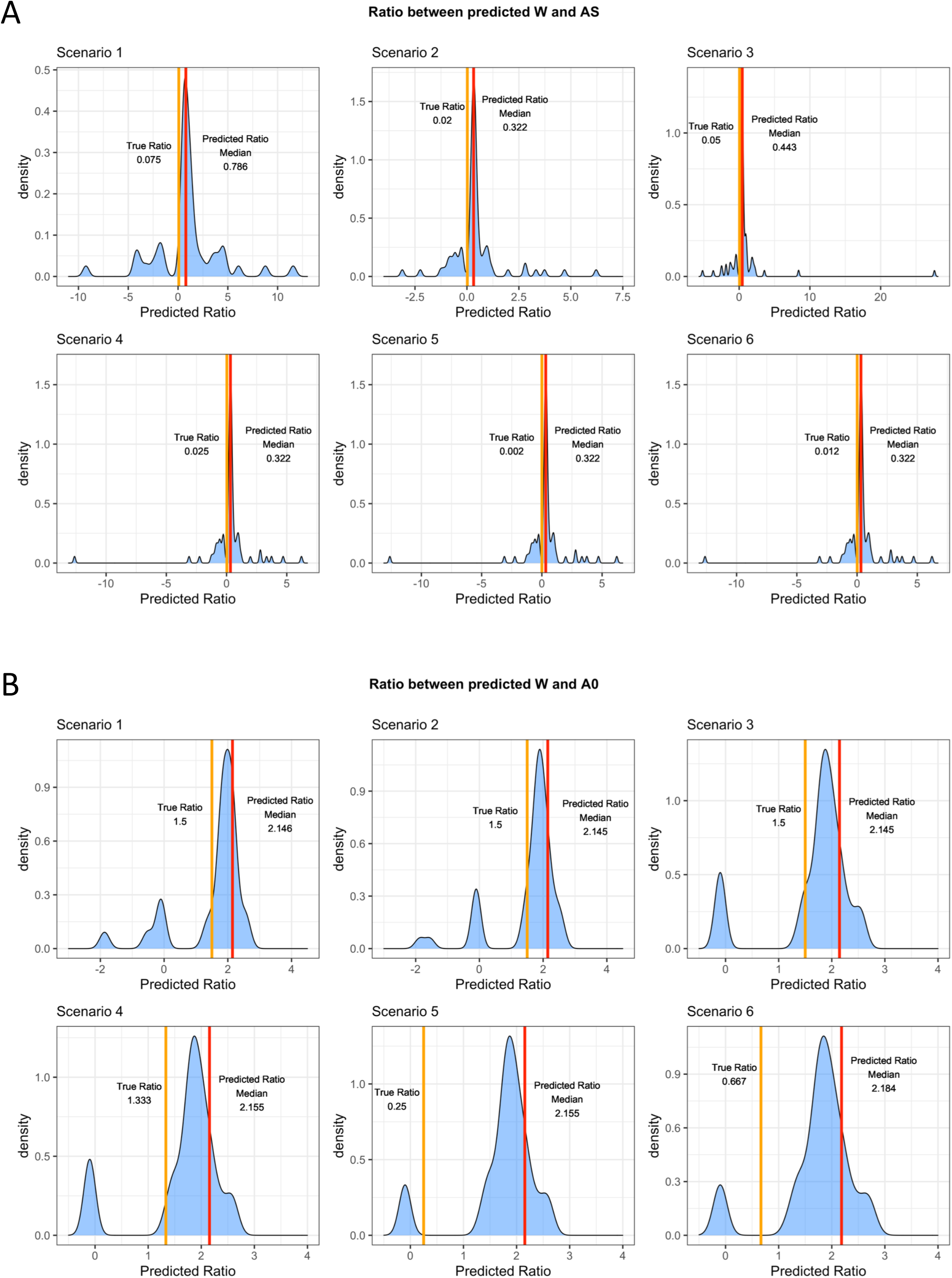
Penalization powers in optimization. A) The distribution of ratios between estimated *W* and the input, *AS*. The red vertical line is the median of the distribution. The orange vertical line is the true ratio in the rewritten optimization algorithm. B) The distribution of ratios between estimated *W* and the input, *A*_0_. The red vertical line is the median of the distribution. The orange vertical line is the true ratio in the rewritten optimization algorithm.

**Supp3.**
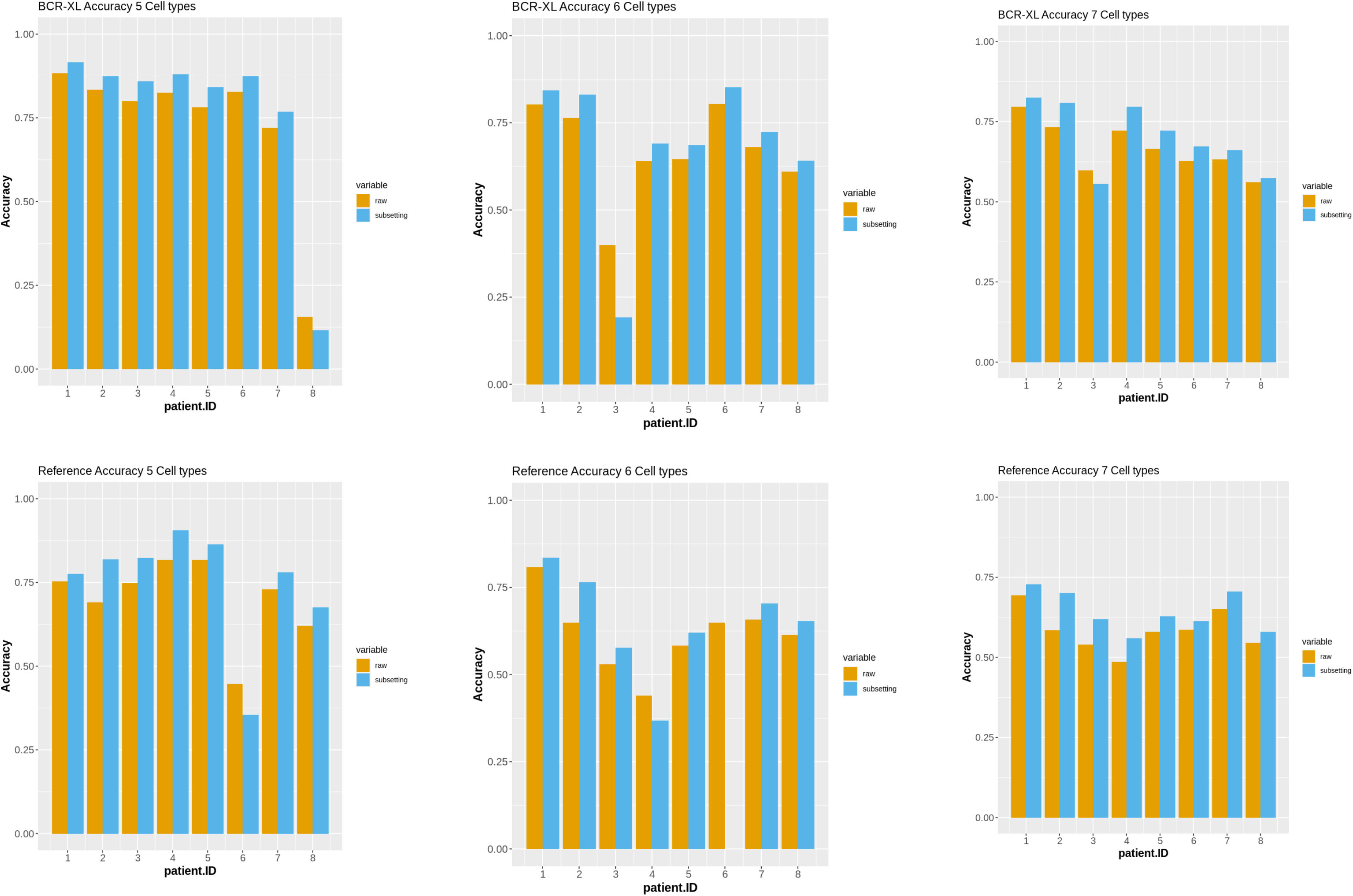
The prediction power on BCR data. The annotation accuracy barplots across samples from stimulated (first row) and unstimulated (second row) groups. The columns are the cell type numbers used (5 cell types in the first column, 6 cell types in the second column, and 7 cell types in the third column). The color represents the raw output or subsetted output from ProtAnno.

**Supp4.**
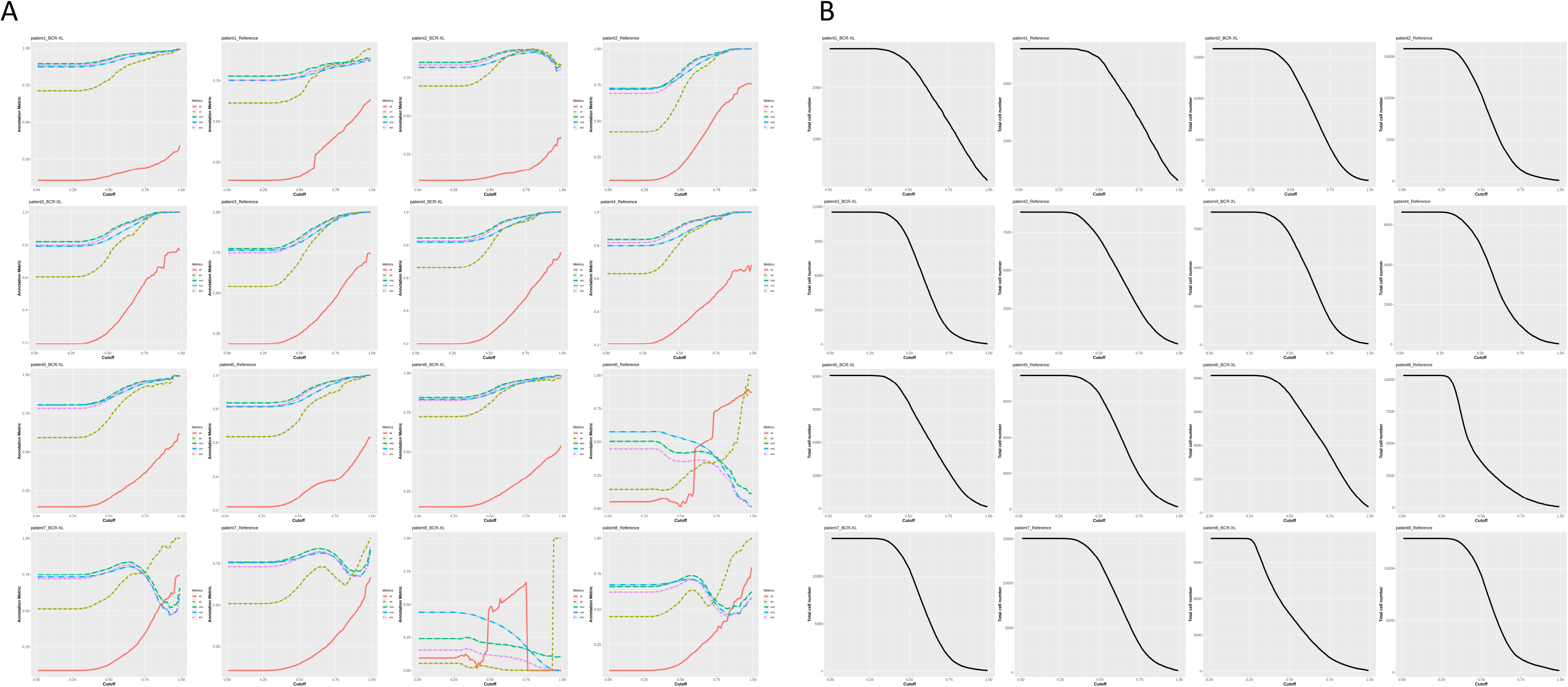
Annotation accuracy improvement by subsetting. A) The annotation metrics with different subsetting cutoffs. The x-axis represents the cutoff value. The y-axis represents the annotation metric value. B) The numbers of kept cells after subsetting filter. The x-axis represents the cutoff value. The y-axis represents the remaining cell counts.

**Supp5.**
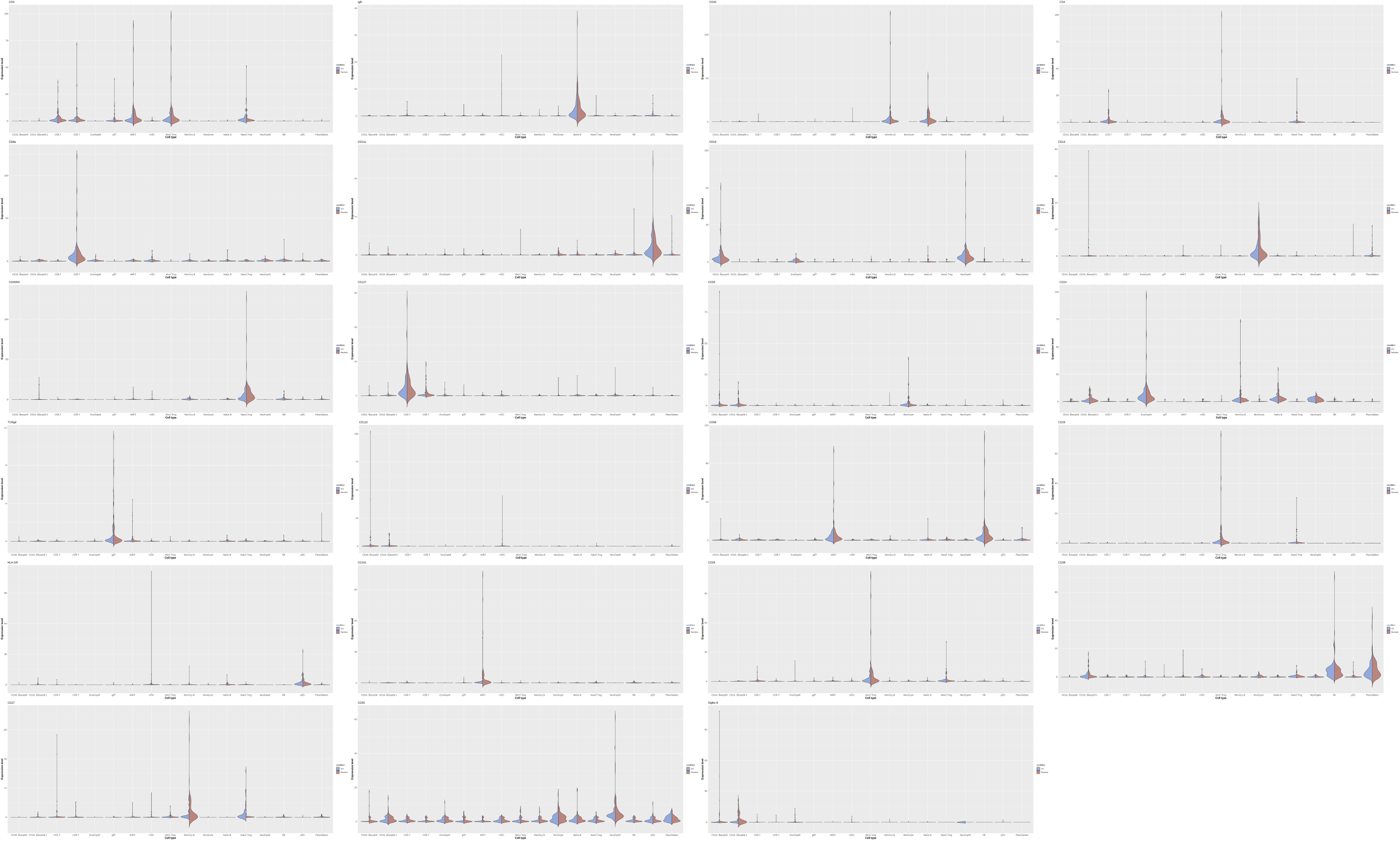
The violin plots of estimated signature expression in in a longitudinal covid-19 study by ProtAnno. The x-axis represents the cell type and the y-axis is the distribution of values in the signature matrix across patients within groups. The colors represent patient groups (ICU and recovery).

**Supp Table 1.**

| Scenario<br>Number | #cell type | #surface<br>markers | #single cell<br>Proteomics<br>cell number | #RNA gene | big_w_mean | big_tau_w | small_w_me<br>an | small_tau_w | p.0 | q.0 | p.neg1 | q.neg1 | mean_var_r<br>atio | corr | cell type<br>proportions<br>ratio |
| --- | --- | --- | --- | --- | --- | --- | --- | --- | --- | --- | --- | --- | --- | --- | --- |
| 1 | 5 | 10 | 2000 | 100 | 2 | 10 | 0.5 | 10 | 0.1 | 0.2 | 0.05 | 0.1 | 10 | 0.5 | 1,2,2,3,3 |
| 2 | 8 | 10 | 2000 | 100 | 2 | 8 | 1 | 8 | 0.15 | 0.3 | 0.1 | 0.15 | 5 | 0.2 | 3,1,2,3,3,1,2,<br>1 |
| 3 | 8 | 10 | 2000 | 100 | 2 | 8 | 1 | 8 | 0.15 | 0.3 | 0.1 | 0.15 | 3 | 0.3 | 3,1,2,3,3,1,2,<br>2 |
| 4 | 8 | 10 | 2000 | 100 | 2 | 7 | 1 | 7 | 0.15 | 0.3 | 0.1 | 0.15 | 3 | 0.2 | 3,1,2,3,3,1,2,<br>3 |
| 5 | 8 | 10 | 2000 | 100 | 2 | 7 | 1 | 7 | 0.2 | 0.4 | 0.1 | 0.2 | 3 | 0.2 | 3,1,2,3,3,1,2,<br>4 |
| 6 | 8 | 10 | 2000 | 100 | 2 | 5 | 1 | 5 | 0.2 | 0.4 | 0.1 | 0.2 | 2 | 0.2 | 3,1,2,3,3,1,2,<br>5 |

## Materials and Methods

### Optimization Procedure

#### Step 1: Update *w* with Lagrangian Multiplier

In this step, we update the k-dimensional non-negative row vector *w_i_*. Thus we have the new loss function by adding the Lagrangian multiplier as equation [eq:updateWlossfunction]. Here to discriminate the primary constraint optimization with the dual Lagrangian function, we use *E* to denote the latter.

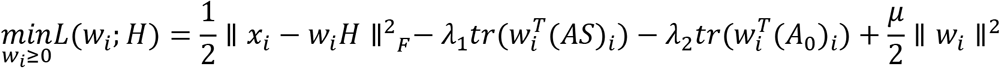

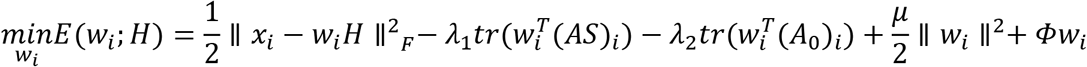

Here *x_i_* and *w_i_* are the ith row of *X* and *W*, respectively. To minimize the loss function by updating row *w_i_*, we have

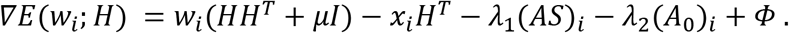

By using the property of *Φw_i_* = 0, we have

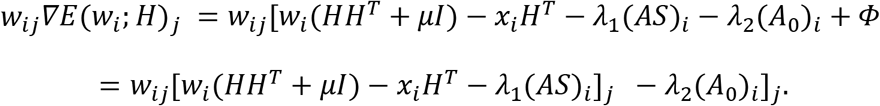

We set the derivative to be zero. To get the non-negative minimizer, we decompose the items that are not non-negative in [eq:2] into positive and negative parts as:

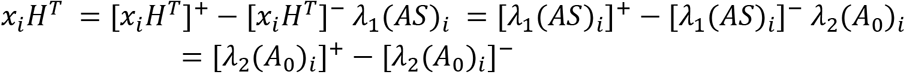

By simple algebra, we have the following equation

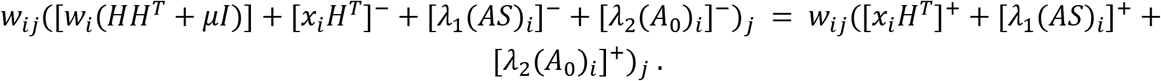

Thus, we update *w_i_* by

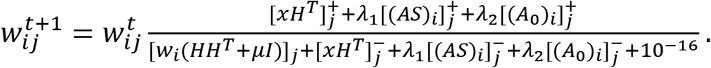

#### Step 2: Update *H* with multiplicative update algorithm

In this step, we optimize *h* column-wise.

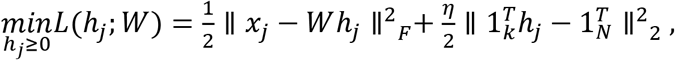

where *W* is the constant matrix and *x_j_* is the *j*-th column vector of *X*. Similarly, by adding the Lagrangian multiplier, we have the following equations.

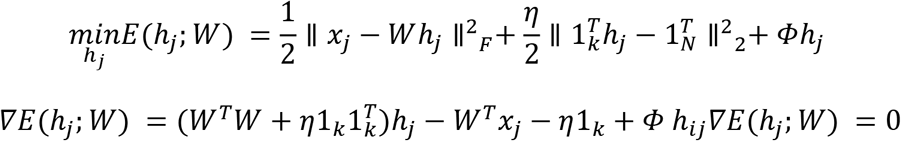

We can update by

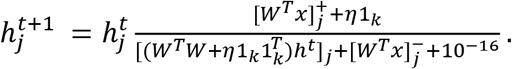

### Simulation Details

The expert matrix *A*_0_ was generated from a multinomial distribution containing three categories, +1, −1, and 0. We generated 100 matrices *A*_0_ and selected the optimal one that had the smallest inner product between the vectors by minimax. Such construction can more likely enable the expression profile X to be a nonnegative linear combination of basis vectors.

In most cases, the expert matrix may be biased. We therefore generated an intermediate discrete matrix 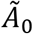 with three possible discrete values, 2, 1, and 0, to represent the expert knowledge matrix used. They were matched with three biomarker distributions within a cell population: high, low, and no expression. In practice, if the entry in *A*_0_ was 1, the corresponding gene would have high expression levels in real data from our observations. Thus, the corresponding element in 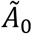 kept the value, 1. However, if the entry in *A*_0_ was −1, it is possible to have low or medium expression level instead of no expression completely. In some rare instances, the corresponding gene might even have high expression. Therefore, the values of −1 would random walk to 1 with probability *q*_−l_ and to 2 with probability *p*_−l_ in 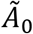. If the entry in *A*_0_ was 0, then the distribution was uncertain. In this case, the values of 0 would random walk to 1 with probability *q*_0_ and to 2 with probability *p*_0_ in 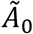. This generated the new discrete matrix 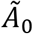 based on the above random walk protocols to distinguish the different biomarkers behaviors.

Next, the signature matrix *W* was generated based on 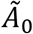 from two positive truncated normal distributions with different expectations. When the entry in 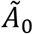 was 2, the corresponding signature gene average expression was sampled from a positive truncated normal distribution with a large mean; when the entry in 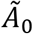 was 1, the corresponding signature gene average expression was sampled from a half-normal distribution with a small mean; otherwise, it was set to 0.1.

After randomly simulating the cell type labels for each single cell in the proteomics expression profile *X*, the expression level was sampled from normal distributions with the expectation of average signature gene expression in *W* and variance calculated from sampled mean and the specified mean-variance ratio.

### Annotation metrics

ARI was calculated using the function adjustedRandIndex from the R package mclust(Scrucca et al. 2016). For NMI, the ground truth and the predicted label for each cell were converted into one-hot vectors and the function NMI in the R package aricode(Vinh, Epps, and Bailey 2009) was used to calculate the NMI for each cell. The reported result was the average across all the cells. Cosine similarity was calculated by first computing the cosine measure between the ground truth one-hot vector and the cluster assignment vector using the cosine function from the lsa package in R, and finally averaging over all the cells. ASW was calculated using the batch_sil function in the R package kBET(Büttner et al. 2018).

### Real Datasets and processing

The BCR CyTOF dataset was downloaded through the HDCytoData(Weber and Soneson 2019) R package. The cell types were labeled by Nowicka et al. (2017)(Nowicka et al. 2017) and could be accessed by the HDCytoData package. The CITE-seq data in Wang et al. (2020)(X. Wang et al. 2020) were downloaded from GEO with accession ID GSE148665. The longitudinal covid-19 CyTOF data were downloaded from (https://brodinlab.com/data-repository/). The two dictionary CITE-seq datasets in Unterman et al. (2020)(Unterman et al. 2020) and Ramaswamy et al. (2020)(Ramaswamy et al. 2021) would be available after they publish the data. The CITE-seq data in Wang et al. (2020)(X. Wang et al. 2020) and Ramaswamy et al. (2020)(Ramaswamy et al. 2021) were manually labeled after the preliminary results by SingleR(Aran et al. 2019). The expert-guided matrices were obtained from the biomarker panels in Unterman et al. (2020)(Unterman et al. 2020).

All the proteomics data were normalized by arcsinh function with cofactor 5. We cleaned and gated the longitudinal CyTOF data by the R package flowCore to filter the intact cells. When processing the association matrix from CITE-seq and transcriptomics signature matrix, all the scRNA-seq data are denoised by SAVERX(J. Wang et al. 2019). The transcriptomics gene list for each application contains the matched protein-coding transcript genes (Table 4) and the top highly variable genes generated by the Seurat(Hao et al. 2020) package.

## Supplementary Materials

### 1 Parameter Tuning

To get the optimal penalty parameters *λ*_1_,*λ*_2_,*μ* and *η*, we considered both the KarushKuhn-Tucker (KKT) condition combined with empirical screening. We first obtained the initial *W* and *H* by setting the arbitrary penalization: *λ*_1_ = 1*, λ*_2_ = 10*, μ* = 50 and *η* = 50. This setting can give an acceptable result in most cases based on our empirical experiments. The first order derivatives of loss function are

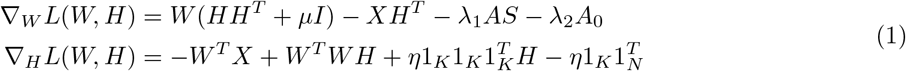

We first initialized *η* by satisfying KKT conditions.

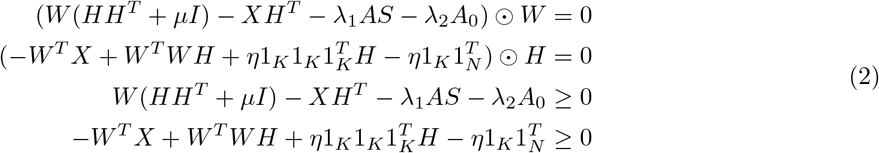

by the second sufficient inequality, we set

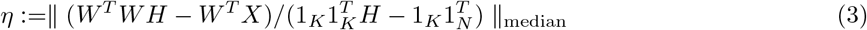

After getting the initial *η*, we initialized *λ*2 and *λ*_1_ in order by minimizing Adjusted Rank Index (ARI) with Louvain clustering. We considered the tuning parameters *λ*2 and *λ*_1_ from 0.1, 1, 10, and 100. The parameter *μ* is charging of the scale and penalization power on signature matrix *W*. Thus, a smaller *μ* can result in a larger norm of *W*. Therefore, we developed a new metric to evaluate the reliability by the difference between the mean value of expression profile *X* and the mean value of signature matrix *W*. To eliminate the non-Gaussian effects, we also considered the difference between the median values of *X* and *W*. Thus, the new metric is formulated as

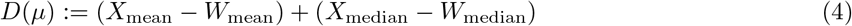

### 2 Theoretical Proofs

#### Convergence of Algorithms

##### Lemma 2.1.

*For any symmetric nonnegative matrix Q* ∈ ℝ^*K×K*^ *and row vector* 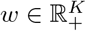 *and* 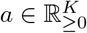*, the following matrix*

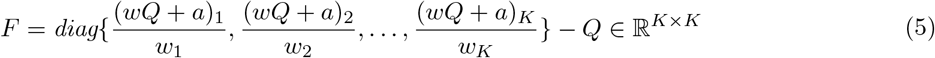

*is always semi-positive definite.*

*Proof.* We construct a new matrix *S* ∈ ℝ^*K×K*^ by *S_ij_* = *w_i_F_ij_w_j_*, where *S_ij_* is the element of *S* whose row is *i* and column is *j*. And we reformulate *F* to be

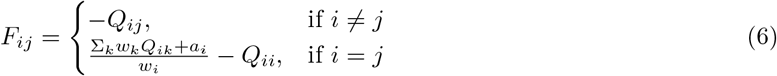

For any nonnegative row vector *v* ∈ ℝ^*K×K*^, we have

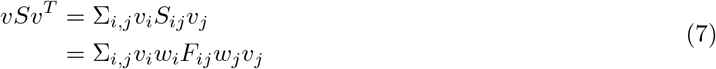

Since row vectors *w* and *v* are nonnegative, *F* is semi-positive definite when *S* is semi-positive definite. Therefore, it is sufficient if we can prove *S* is semi-positive definite. In the following, we follow equation 7 and prove that the product is always nonnegative.

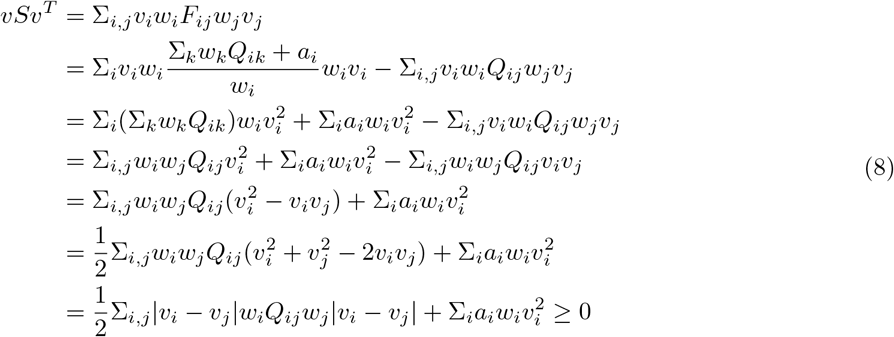

The first term in the above formula is nonnegative since *w* is nonnegative and *Q* is semi-positive definite. The second term is always greater than or equal to 0 due the nonnegativity of *a* and *w*. Thus, *S* is a semi-positive definite matrix.

##### Theorem 2.2.

*Consider the following quadratic optimization problem,*

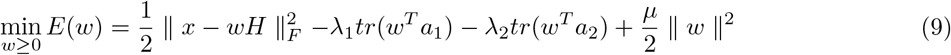

*which is the loss function for a row vector w* ∈ ℝ^*K*^. *In this optimization problem, H* ∈ ℝ^*K×N*^ *is a constant non-negative matrix, and x, a*_1_, *a*_2_ ∈ ℝ^*K*^ *are constant row vectors. The penalty parameters, λ_1_, λ_2_, and μ are positive numbers. The the following update rule*

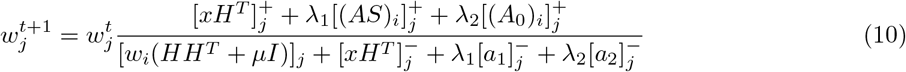

*where we denote*

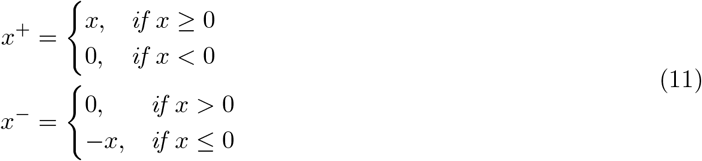

*converges to its optimal solution with the convergence rate*

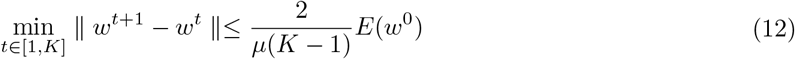

*Proof.* To prove that there exists a *w^t^*^+1^ that *E*(*w^t^*^+1^) ≥ *E*(*w^t^*), we would like to construct an auxiliary function *F* (*w, w^t^*), s.t.

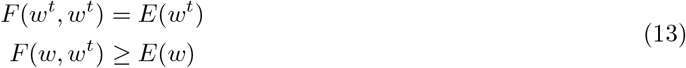

Then we have

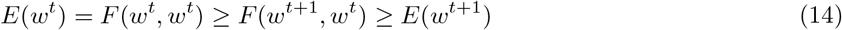

where *w^t^*^+1^ = arg min_*w*_ *F* (*w, w^t^*). Thus, the loss function *E*(*w^t^*) is monotonously non-increasing w.r.t the iteration *t*. We define the auxiliary function as the following.

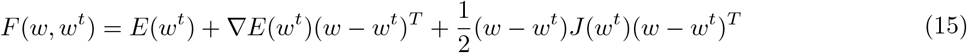

where

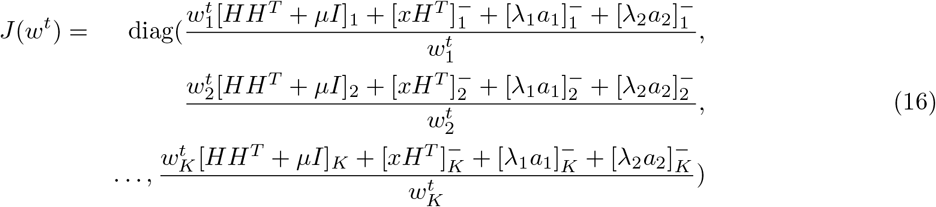

We can approximate *E*(*w^t^*) based on Taylor expansion.

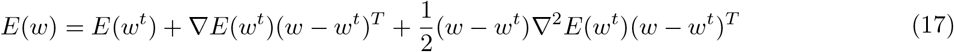

where, by simple derivation,

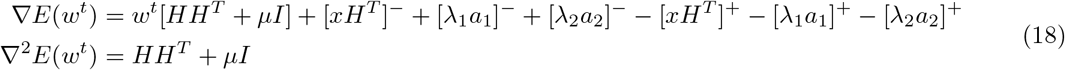

Now we can have 14 satisfied if *F* (*w^t^*^+1^*, w^t^*) ≥ *E*(*w^t^*^+1^). To get it, we have

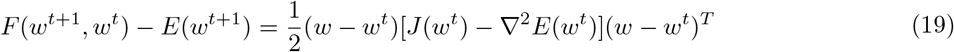

Obviously, ▽^2^*E*(*w^t^*) = *HH^T^* + *μI* is a positive definite matrix since *μ* is positive. Furthermore, the row vector

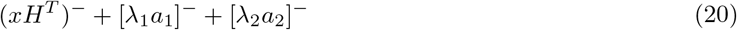

is always nonnegative. Due to the 2.1, *J* (*w^t^*) − ∇^2^*E*(*w^t^*) is a semi-positive definite matrix. So that *F* (*w^t^*^+1^*, w^t^*) ≥ *E*(*w^t^*^+1^) is always satisfied and *E*(*w^t^*) is nonincreasing w.r.t to *t*. Next, we would like to find the optimizer of *F* (*w, w^t^*). To achieve this, we set 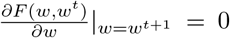, which is equivalent to equation ∇*E*(*w^t^*) + (*w^t^*^+1^ − *w^t^*)*J* (*w^t^*) = 0. Therefore, we have

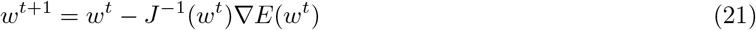

For each element of *w^t^*^+1^, the above formula can be reformulated as

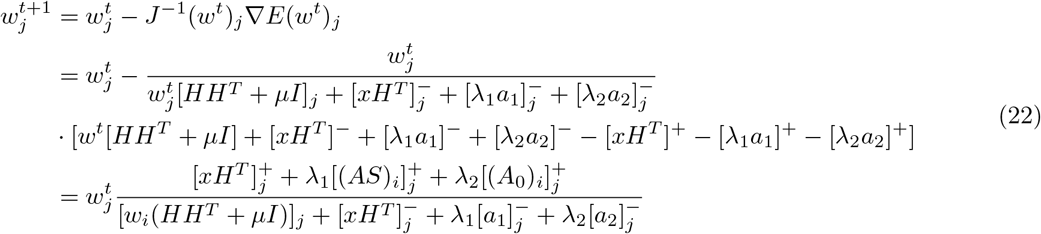

So far we have proved that the above updating rule can make *E*(*w^t^*) nonincreasing w.r.t to the *t*. Specifically, we have the lower bound for every element on the diagonal of *J* matrix.

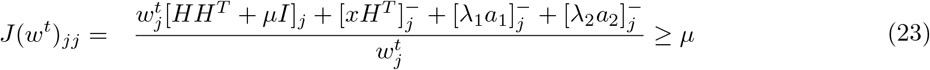

Based on 14 and 23, we can derive the lower bound of the difference between two loss function values after one update.

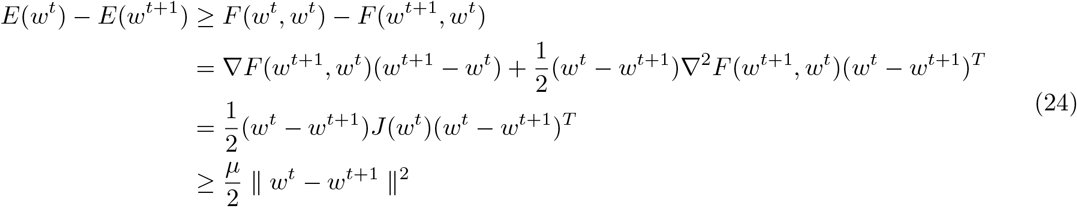

since ▽*F* (*w^t^*^+1^*, w^t^*) = 0 and ▽^2^*F* (*w^t^*^+1^*, w^t^*) = *J* (*w^t^*). And by accumulating ‖*w^t^ − w^t^*^+1^‖^2^ from 0 to *T* − 1, we then can have

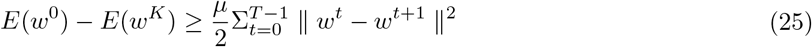

Thus, for any given fixed iteration number *T*, we can have

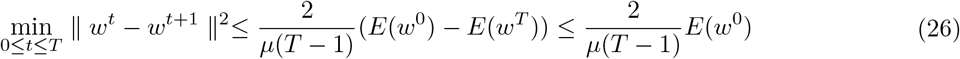

##### Theorem 2.3.

*Consider the following quadratic optimization problem,*

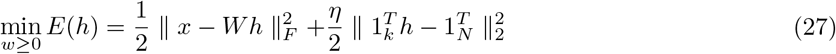

*which is the loss function for a column vector h* ∈ ℝ^*K*^. *In this optimization problem, W* ∈ ℝ^*D×K*^ *is the constant non-negative matrix, and x* ∈ ℝ^*K*^ *is a constant column vectors. The penalty parameter, η is a positive number. The the following update rule*

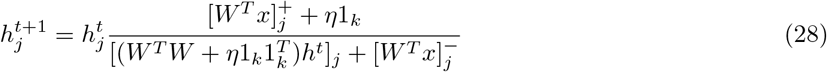

*converges to its optimal solution with the convergence rate*

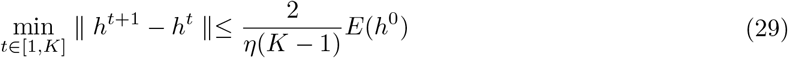

*Proof.* To prove that there exists an *h^t^*^+1^ that *E*(*h^t^*^+1^) ≥ *E*(*h^t^*), we can construct an auxiliary function *F* (*h, h^t^*), s.t.

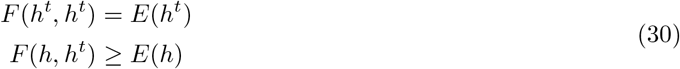

Then we have

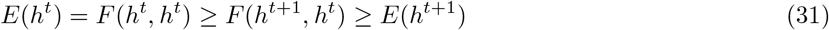

where *h^t^*^+1^ = arg min_*h*_ *F* (*h, h^t^*). Thus, the loss function *E*(*h^t^*) is monotonously non-increasing w.r.t the iteration *t*. We define the auxiliary function as the following.

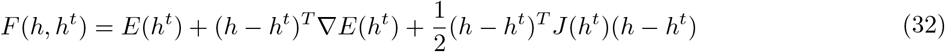

where

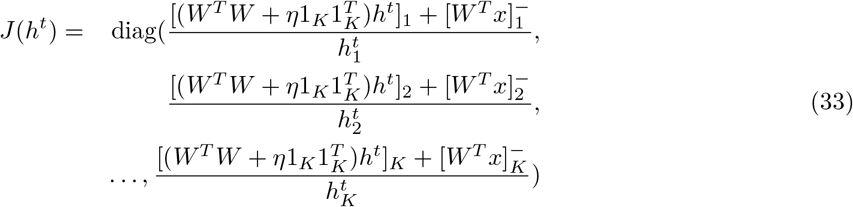

We can approximate *E*(*h^t^*) based on Taylor expansion.

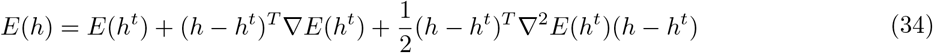

where, by simple derivation,

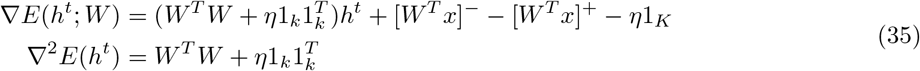

Now we can have 31 satisfied if *F* (*h^t^*^+1^*, h^t^*) ≥ *E*(*h^t^*^+1^). To get it, we have

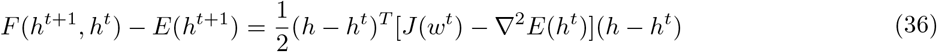

Obviously, 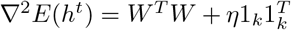 is a positive definite matrix since *η* is positive. Furthermore, the row vector

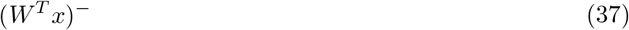

is always nonnegative. Due to the 2.1, *J* (*h^t^*) − ∇^2^*E*(*h^t^*) is a semi-positive definite matrix. So that *F* (*h^t^*^+1^*, h^t^*) ≥ *E*(*h^t^*^+1^) is always satisfied and *E*(*h^t^*) is nonincreasing w.r.t to *t*. Nextly, we would like to find the optimizer of *F* (*h, h^t^*). To achieve this, we set 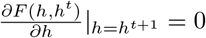, which is equivalent to equation ∇*E*(*h^t^*) + (*h^t^*^+1^ − *h^t^*)*J* (*h^t^*) = 0. Therefore, we have

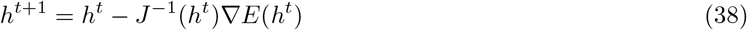

For each element of *h^t^*^+1^, the above formula can be reformulated as

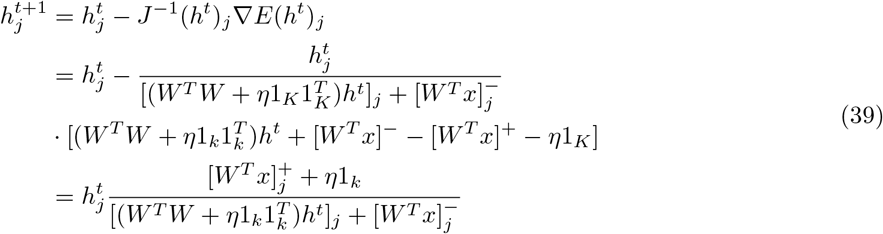

So far we have proved that the above updating rule can make *E*(*h^t^*) nonincreasing w.r.t to the *t*. Specifically, we have the lower bound for every element on the diagonal of *J* matrix.

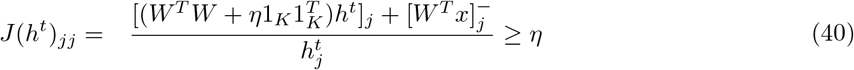

Based on 31 and 40, we can derive the lower bound of the difference between two lose function values after one update.

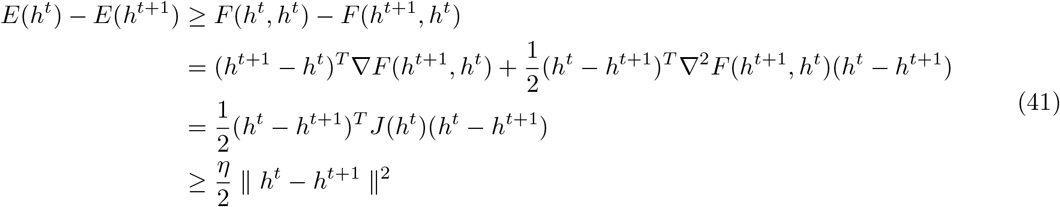

since ▽*F* (*h^t^*^+1^*, h^t^*) = 0 and ▽^2^*F* (*h^t^*^+1^*, h^t^*) = *J* (*h^t^*). And by accumulating ‖*h^t^ − h^t^*^+1^‖^2^ from 0 to *T* − 1, we have

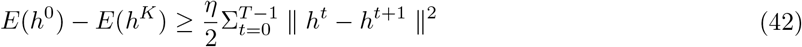

Thus, for any given fixed iteration number *T*, we have

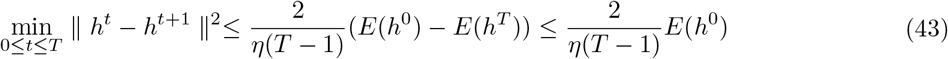

### 3 Rewriting the Loss Function

To conduct the parameter analysis and validate the optimization convergence, we rewrote the original loss function to a equivalent formula as following.

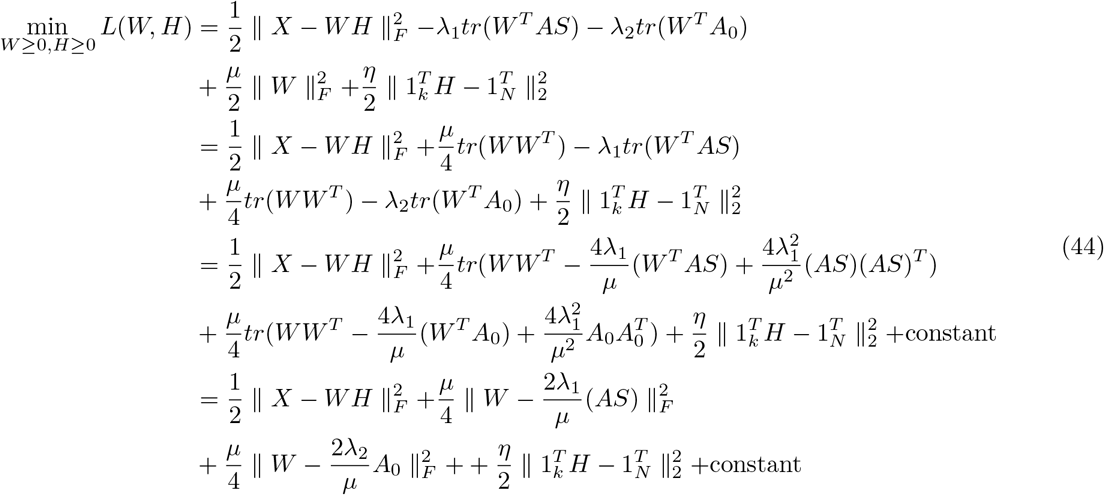

## Notes

### Competing Interest Statement

The authors have declared no competing interest.

